# USP1 acts as a chaperone for HSFA2 and plays a crucial role in thermopriming in Arabidopsis

**DOI:** 10.1101/2025.01.21.634064

**Authors:** Prabhu Manickam, Kirti Shekhawat, Anam Fatima, Hanna M. Alhoraibi, Aala A. Abulfaraj, Naganand Rayapuram, Heribert Hirt

## Abstract

Plants employ diverse strategies to cope with different types of heat stress. The response to short-term acute heat stress differs significantly from that to moderate heat stress followed by severe stress events. After experiencing moderate heat stress, plants exhibit a more robust response to subsequent severe stress, a phenomenon known as thermopriming or acquired thermotolerance. Thermopriming creates a memory by maintaining the heat stress (HS) memory-related genes in an alert state. In this work, we investigated the role of Arabidopsis Universal Stress Protein 1 (USP1) in plant heat stress responses. CRISPR-Cas9 generated knockout *usp1* mutant lines showed no morphological changes during development and normal growth conditions. However, *usp1* mutant plants showed enhanced levels of apoplast hydrogen peroxide and superoxide reactive oxygen species accumulation upon heat stress. Transcriptome analyses revealed that genes related to protein folding, electron transport, and oxidative phosphorylation are strongly upregulated in *usp1* mutant plants. USP1 is essential for acquired thermotolerance, as *usp1* mutants are compromised in heat stress memory but show normal responses to acute heat stress similar to *hsfa2* mutants. Biochemical assays showed that USP1 functions as a molecular chaperone, protecting the transcription factor HSFA2 from heat-induced denaturation. Moreover, *usp1* mutant plants show decreased transcript levels of heat stress response genes and reduced H3K4me3 enrichment at memory gene loci. These data show that USP1 plays an important role as a chaperone of HSFA2 in mediating plant heat stress memory.

## Introduction

Plants are sessile organisms that are constantly exposed to numerous changing environmental conditions. Heat stress (HS) causes a broad range of irreversible cellular damages, such as modifying the membrane permeability, altering the protein stability, denaturation, aggregation of intracellular proteins, and disrupting the metabolic processes (Wu et al., 2019). Plants have developed sophisticated mechanisms to enhance their survival adaptability to temperatures above the optimum level for their growth and development. Heat stress has a complex impact on cellular function, which involves complex mechanisms and processes that contribute to thermotolerance. Some of them are integral to basal thermotolerance, and others are activated during acquired thermotolerance. The acquisition of thermotolerance relies on the activation of the heat stress response (HSR) to maintain protein stability, minimize cellular damage, and adjust metabolism. During heat stress, heat shock transcription factors (HSFs) regulate the expression of heat shock proteins (HSPs), which act as chaperones to protect and stabilize proteins. Among the 21 HSFs in *Arabidopsis thaliana* (hereafter, *Arabidopsis*), eight are involved in HSR, with HSFA2 being a key player (Charng et al., 2007; Schramm et al., 2008). HSFA2, which is highly induced by heat stress, is crucial for both developing and maintaining thermotolerance (Nishizawa et al., 2006; Schramm et al., 2006; Charng et al., 2007). The inherent ability to survive and recover from acute heat stress is called basal thermotolerance (Larkindale et al., 2005). Plants utilize acquired thermotolerance to build resilience against subsequent episodes of more intense HS (Hong and Vierling et al., 2000; Larkindale et al., 2005). Acquired thermotolerance is supported by epigenetic mechanisms, for example, HSFA2 promotes the accumulation of H3K4 methylation at heat stress memory genes, ensuring their sustained expression and providing continued protection against recurring heat stress conditions.

USPs have been identified in diverse plant species including *Hordeum vulgare*, *Arabidopsis*, *Oryza sativa, Gossypium arboreum, Astragalus sinicus, Vicia faba, Solanum pennellii*, and *Viridiplantae*, however, their functions remain largely unknown. *Arabidopsis* contains 44 proteins that are similar to the *Methanococcus jannaschii* protein family of USPs that contain an ATP binding site but different from the *E.coli or Haemophilus influenza* USP protein family lacking the ATP binding residues (Sousa and Mckay et al., 2001). The rate of cell survival during prolonged exposure to stress agents appears to be increased by USP, and this may give plants a broader spectrum of stress tolerance. For example, a study on the wild tomato *Solanum pennellii* showed that the SpUSP protein is present in both the nucleus and associated with the cell membrane (Loukehaich et al., 2012). Transcript levels of SpUSP increase in response to drought in leaves reducing stomatal aperture to protect plants from drought. There are several reports which showed that USP-like proteins play an important role in transport and scaffolding (Sinha et al., 2016).

High temperatures lead to the overproduction of ROS and oxidative stress. Plants have evolved antioxidant mechanisms to alleviate the effect of ROS and protect cells from damage (Gill and Tuteja et al., 2010). HSPs act as molecular chaperones by maintaining cellular homeostasis challenged by heat-induced damage by preventing the misfolding, aggregation, degradation, and denaturation of proteins under stress conditions (Hong and Vierling et al., 2000, 2001). AtUSP (At3g53990) exhibits a chaperone function and is significantly induced by heat, H_2_O_2_, and drought treatments (Jung et al., 2015). Overexpression of AtUSP in transgenic plants results in resistance to heat stress (Jung et al., 2015; Melencion et al., 2017). Furthermore, AtUSP has been shown to play a role as an RNA chaperone during cold stress in *Arabidopsis.* RNA molecules tend to misfold at lower temperatures and become inactive. The helper protein AtUSP functions as an RNA chaperone to unwind misfolded molecules back to their folded (native form) so they can provide a template for the translation machinery (Kang et al., 2013; Jung et al., 2015). There are numerous examples of USP proteins functioning as chaperones and redox proteins (Chuang et al., 2006; Park et al., 2009; Chi et al., 2013). These molecular properties suggest that AtUSP functions as a molecular chaperone.

Previous studies suggested that the chaperone function of AtUSP was increased by heat treatment and overexpressor lines of AtUSP show resistance to heat shock and oxidative stress (Jung et al., 2015). In this study we investigated the function of USP1 in heat resistance by correlating the physiological, transcriptomic analysis, and biochemical study. We generated *usp1* loss of function mutants using CRISPR-Cas9 technology. We provide evidence that *usp1* mutants are compromised in heat stress memory but not in the basal heat stress response. We observed that *usp1* mutant plants have an HS memory-specific phenotype, similar to *hsfa2* mutants. We demonstrate that *Arabidopsis usp1-1* and *usp1-2* knockout mutant lines accumulate ROS and express stress-related genes. By differential scanning fluorimetry (DSF), we show that the USP1 protein functions as a chaperone on the key heat stress memory factor HsfA2.

## Materials and Methods

### 1. Plant Materials and Growth Conditions

*Arabidopsis thaliana* ecotype Columbia-0 (Col-0) was used as wild-type (WT) and *usp1-1*, *usp1-2*, and *hsfa2* mutants were in the Col-0 background. Seeds were surface-sterilized and stratified at 4°C on growth medium for at least two days. The growth medium was composed of ½ Murashige and Skoog basal salts with minimal organics, 0.05% MES hydrate, and 0.5% agar type (Sigma A4675, St. Louis, MO, USA). The pH was adjusted to 5.7 with KOH. Seedlings were grown at 23°C day/22°C night in long day conditions, i.e., 16 h light/8 h dark photoperiod, for the indicated time. For growth in Jiffy pots in Percival growth chambers, seeds were stratified at 4 °C in water for at least two days and then grown at 23°C day/22°C night, 60% humidity in short day conditions, i.e., 8h light/16 h dark photoperiod for four weeks.

### 2. Design of guide RNA (gRNA), construct construction, and Agro transformation

To generate *USP1* knockout mutant lines in *Arabidopsis*, we designed 20- nucleotide guide RNAs (gRNAs) specifically targeting the USP1 gene using the CRISPR-P 1.0 tool (Wang ZP et al., 2015). Two gRNAs with the lowest off-target scores were selected. The gRNA expression cassettes were cloned into the mCherry-pHSE401 vector containing an egg cell-specific promoter, as previously described (Wang ZP et al., 2015, Table 1). These constructs were then transformed into Arabidopsis Col-0 plants via the Agrobacterium-mediated floral dip method. Seeds harvested from T0 plants were screened on ½ MS media plates containing hygromycin to identify resistant seedlings, which were then transplanted into soil. Genomic DNA was extracted from T1 transgenic lines for sequence analysis. To detect mutations in *USP1*, PCR was used to amplify DNA fragments flanking the gRNA target sites, with *USP1*-specific primers. PCR products were then purified, sequenced and analyzed using CLC Workbench to identify CRISPR-Cas9- induced insertions/deletions (Figure S1). Counterselection PCR using Cas9-specific primers was performed to screen for non-transgenic T2 plants, confirming that the desired mutations were not linked to the transgene (Figure S2A–C, Table 2). This process enabled the identification of USP1 knockout mutant lines in Arabidopsis.

**Table 1.**
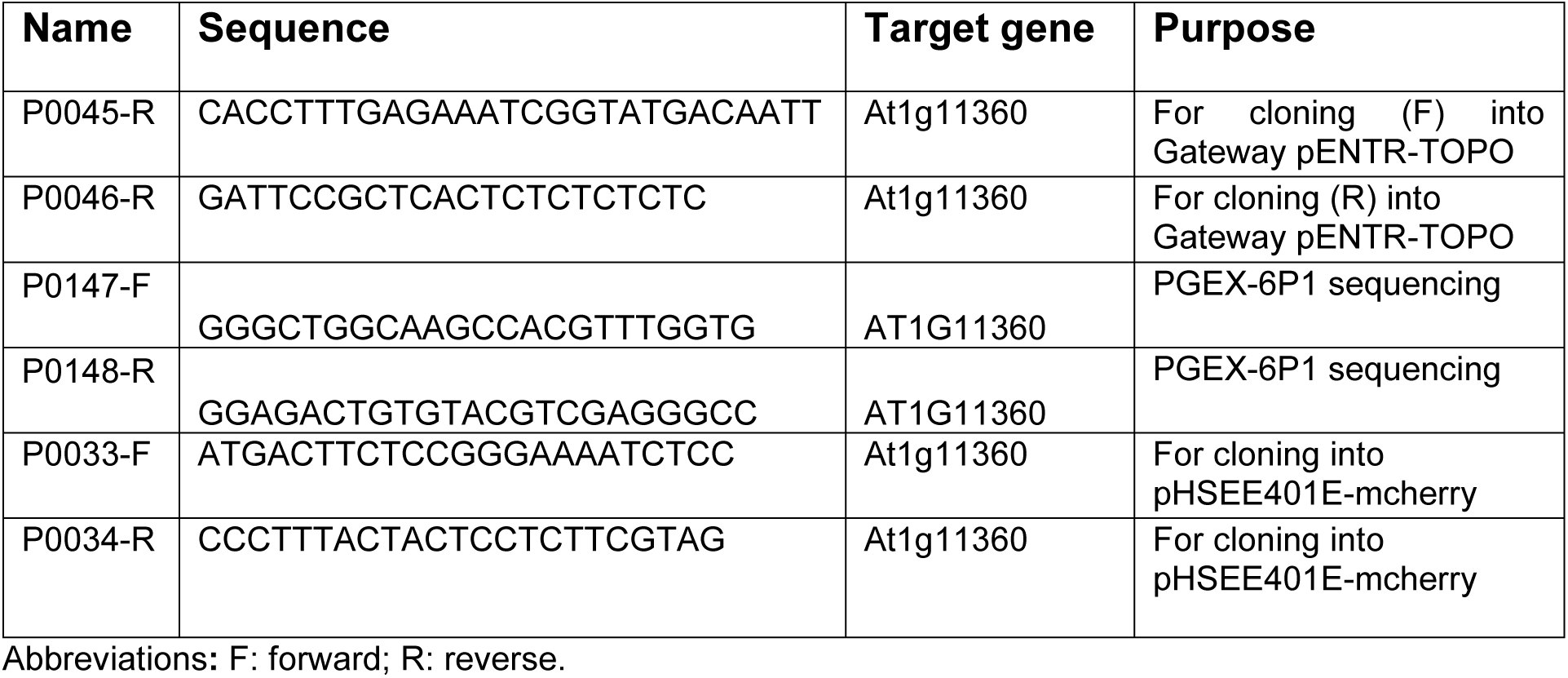
Primers Used For Cloning.

**Table 2.**
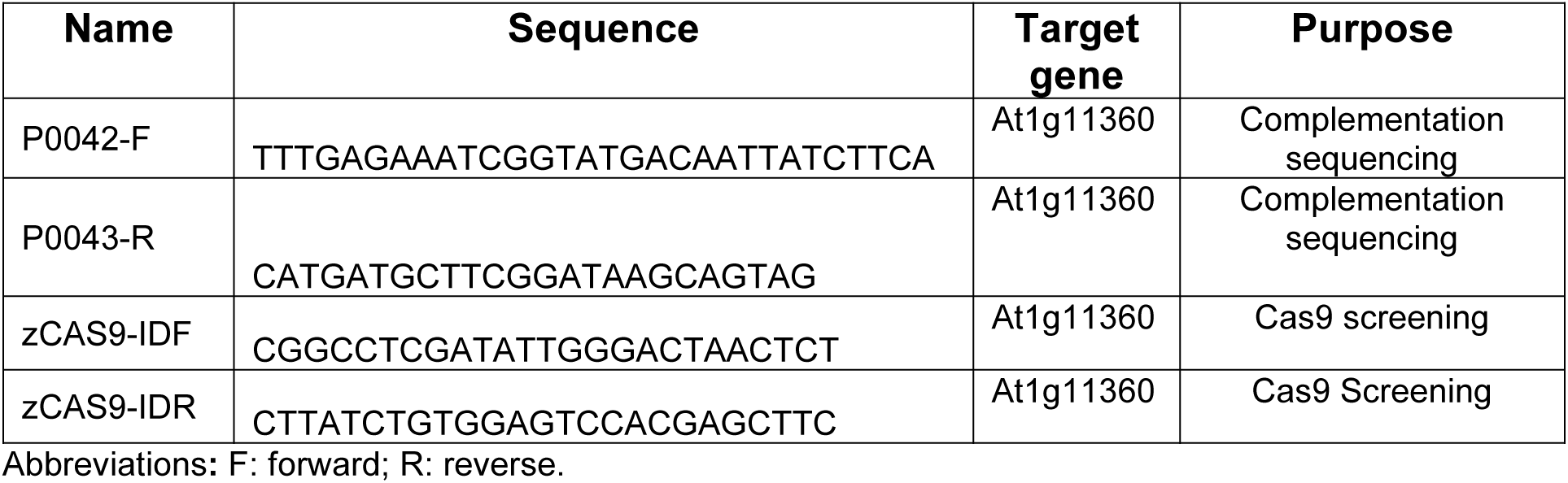
Primers Used For Sequencing.

### 3. Quantitative Realtime-PCR

WT and mutant *Arabidopsis* seedlings were grown on ½ MS agar for 14 days. The samples were then homogenized in liquid nitrogen, and total RNA was extracted using the NucleoSpin Plant RNA kit. 1 µg of total RNA from each sample was reverse transcribed into cDNA using the SuperScript III First-Strand Synthesis SupermixKit (Invitrogen, Vilnius, Lithuania). The resulting cDNA was used as a template for qRT-PCR using the heat stress-responsive genes primers (Table 3) and SsoAdvanced Universal SYBR Green Supermix (Bio-Rad, Germany). ACTIN (At3g18780) and UBQ10 (At4g05320) were amplified as internal control genes to normalize gene expression levels in all samples using BioRad CFX software v 3.0. The normalized gene expression levels were expressed relative to wild-type controls in each experiment.

**Table 3.**
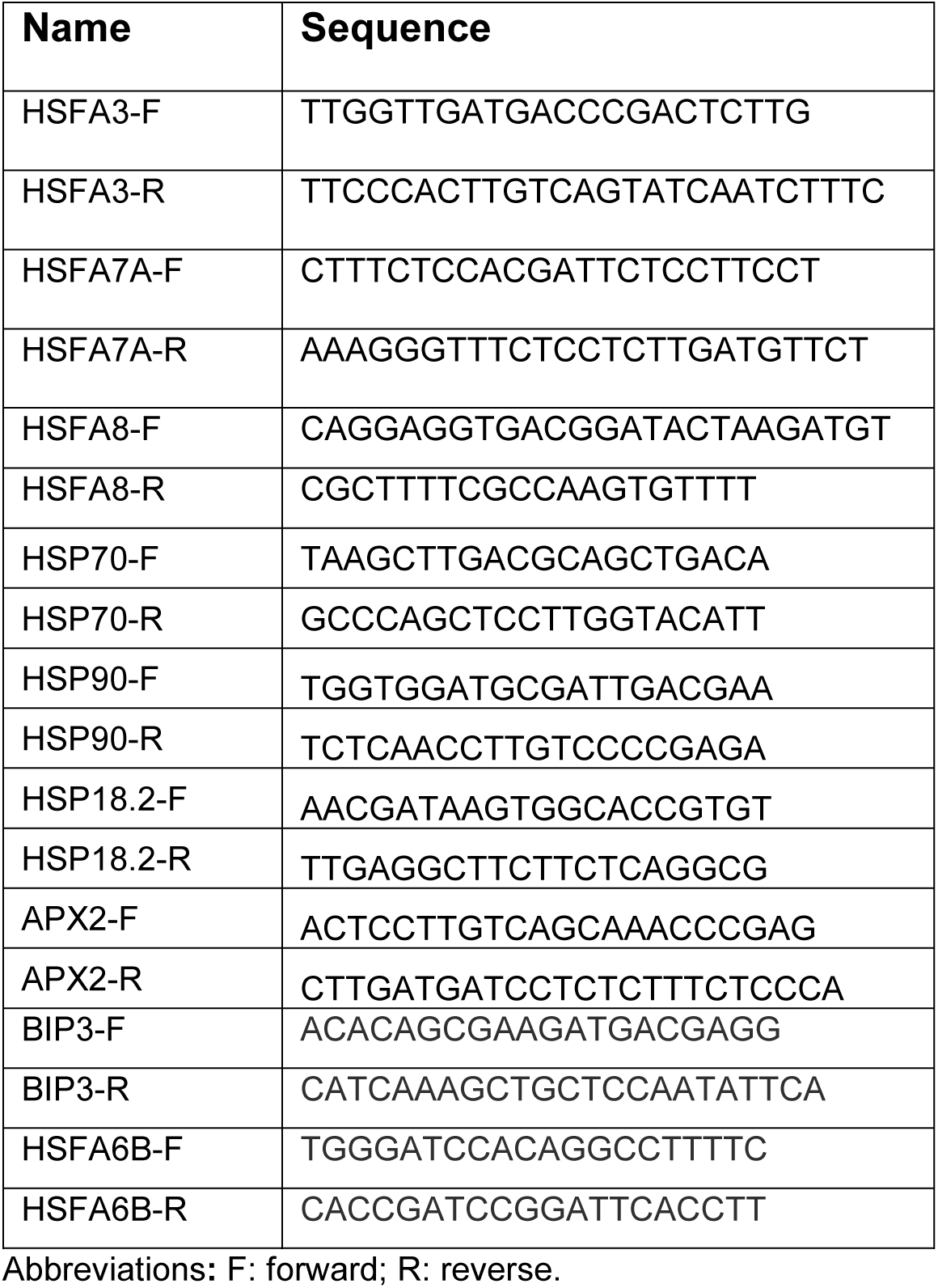
Primers Used For qPCR.

### 1. 3. *In situ* detection of H_2_O_2_ and O_2_^-^

To visualize the superoxide O_2_^-^ and hydrogen peroxide H_2_O_2_ levels in fully developed leaves from *Arabidopsis* plants grown for four weeks in jiffy pots was performed with minor modifications as described by Daudi, A. and O’Brien, J. A. (2012). The four weeks old leaves were stained with nitroblue tetrazolium (NBT) (N6876, Sigma-Aldrich, Manheim, Germany) and 3,3′-diaminobenzidine (DAB) (D5637, Sigma-Aldrich, St. Louis, MO, USA), respectively. The leaves were then placed on slides that contained 50% glycerol. The pictures were captured using a Nikon SMZ25 stereomicroscope.

## 4. RNA Sequencing

Three independent biological replicates of 14-day-old WT and mutant *Arabidopsis* seedlings were grown on ½ MS agar. RNA seq libraries were prepared using the Illumina Truseq Stranded mRNA sample Preparation LS (low sample) kit according to the manufacturer’s protocol and sequenced on a HiSeq 4000 platform with a read length of 300 bp paired ends. Reads were quality controlled using FASTQC v0.11.5 (http://www.bioinformatics.babraham.ac.uk/projects/fastqc/). Trimmometric was used for quality trimming. The read quality filter parameters were set as follows: minimum length of 36 bp; mean Phred quality score greater than 30; remove leading and trailing bases with the base quality below 3; sliding window of 4:15. TopHat v2.1.1 was used to align short reads to the *Arabidopsis thaliana* TAIR 10 genome and for transcript assembly, and to measure differential expression levels (fragment per kilobase of transcript per million mapped reads, FPKM). The differentially expressed genes (DEGs) were identified using Cufflinks and the limma package in R. Cuffdiff v2.2.1 (Trapnell et al., 2012) with quartile normalization. Genes with a two-fold change between samples with and without flg22 treatment and P-values ≤0.05 were considered significantly differentially expressed. Hierarchical clustering of these genes was performed using Gene ontology (GO) classification was performed with agriGo v2 (Du, Zhou, et al., 2010) and DAVID Bioinformatic Resources 6.8 v2022q1 (https://david.ncifcrf.gov/, Dennis, et al., 2003). Enrichment in GO terms was calculated and adjusted with a p-value of 0.05. The minimum number of mapping entries for a GO category was set to be 5. The enrichment figures were produced using the R package ggplot2. RNA-Seq data are available at NCBI’s Gene Expression Omnibus GEO Series under accession number GSE286899.

## 5. Heat priming and heat stress

*Arabidopsis* WT and mutant lines were grown on the ½ MS plates for five days. Seedlings of the same size were transferred to new ½ MS plates. On the 9^th^ day after transfer, plants were exposed to 37^0^C of heat acclimation for 3 hours, followed by two days of recovery at 22^0^C in Percival incubators. Plants were further treated with 44^0^C heat stress for 30 min in a water bath. The samples were collected at the indicated time points when fresh weight and survival rates were measured. Three technical replicates were tested in three independent experiments.

## 6. Protein purification

The RIKEN full-length cDNA was used to clone the *Arabidopsis* USP1 coding sequence (CDS) by PCR, which was then cloned into the pENTR/D-Topo entry vector in accordance with the manufacturer’s instructions using the pENTR/D-Topo cloning kit (Invitrogen; Cat. No. K2400-20). The resulting fragments were digested with BamHI and XhoI and ligated into the pGEX-6P-1 expression vector (GE Healthcare, NJ, USA; Table 3). Plasmid containing the USP1 coding sequence in pGEX-6P-1 was introduced into *E. coli* BL21 (DE3) cells. Cells were grown in LB broth with ampicillin (100 mg/mL) at 37°C until an OD600 nm of 0.6, and expression was induced by adding a final concentration of 1 mM IPTG, and cultures were incubated at 16°C overnight in a shaker incubator. Cells were harvested by centrifugation at 8,000 g for 10 minutes. A cell pellet from a one-liter culture was resuspended in 25 mL of lysis solution (50 mM Tris-HCl (pH 8.0), 200 mM NaCl, 3 mM DTT, 700 mM Triton X-100, or 9.6 mM Tween-20), and then sonicated to lyse the cells. Proteins were purified using Glutathione Sepharose 4B resins and the resultant supernatant was centrifuged at 30,000 g for 30 min to eliminate cell debris (GE Healthcare). Following GST cleavage, the protein was further purified using a solution containing 10 mM HEPES (pH 7.5), 50 mM KCl, and 3 mM DTT on a HiLoad16/60 Superdex 200 prep-grade gel filtration column (GE Healthcare). The N-terminal GST tag on USP1 was removed by incubating it overnight at 4°C with PreScission Protease (Sigma, Darmstadt, Germany). The purified protein was concentrated to 10 mg/mL and stored at -80°C. Eluted proteins were passed through gel filtration equilibrated with a buffer containing 10 mM HEPES (pH 7.5), 50 mM KCl, and 3 mM DTT and further utilized for *in vitro* experiments.

## 7. Differential Scanning fluorimetry (DSF)

In a 96-well microtiter plate, purified proteins USP1, LDH and HSFA2 of different concentrations 0.5 µM, 1 µM, 5 µM, 10 µM, and 15 µM were added to SYPRO orange (Thermo Fisher Scientific Ref S6650, Lot 2008138) in phosphate buffer, with a final concentration of 1X to a final volume of 10 µL per well (Fig. S3). The Bio-Rad CFX 96 qPCR was used for DSF experiments, which were heated from 25^0^C-100^0^C at a rate of 2^0^C/min. Using Bio-Rad CFX 96 qPCR, the fluorescence was monitored. Experiments were performed in triplicates.

## 8. ChIP-qPCR

We performed chromatin immunoprecipitation (ChIP) based on the method described by Lämke et al. (2016a). Approximately 500 mg of 10- and 12-day-old seedlings were cross-linked by vacuum infiltration with 1% formaldehyde for 15 minutes. The cross-linking was quenched using 2 M glycine, and the samples were stored at −80°C for later processing. Nuclei were extracted, and chromatin was sonicated using a Diagenode Bioruptor (medium setting, 14 cycles of 30 seconds on/off with ice cooling), producing DNA fragments around 250 bp in size. Antibodies (anti-H3, ab1791; anti- H3K4me3, ab8580, both from Abcam) were pre-coupled with protein A-coated agarose beads (Invitrogen) for at least 2 hours at 4°C. Immunoprecipitation was performed overnight at 4°C in IP buffer. After washing and reverse cross-linking, DNA was purified using the phenol-chloroform extraction method, precipitated with ice-cold ethanol and glycogen (Invitrogen), and re-suspended in 20 µL of water. We conducted ChIP-PCR for three regions of the *APX2* and two regions of the *HSP18.2* gene (Table 4). Amplification results were normalized to the H3 signal (signal for the modification normalized to the H3 signal). The values shown in the graphs represent the means of three biological replicates.

**Table 4.**
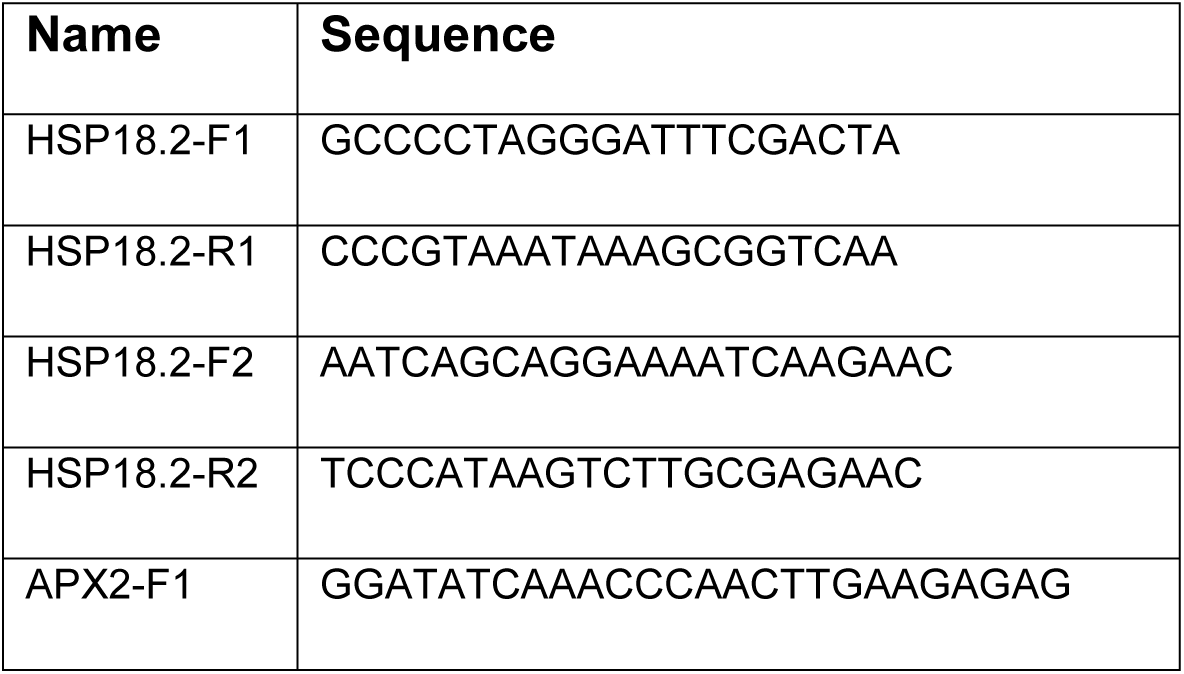
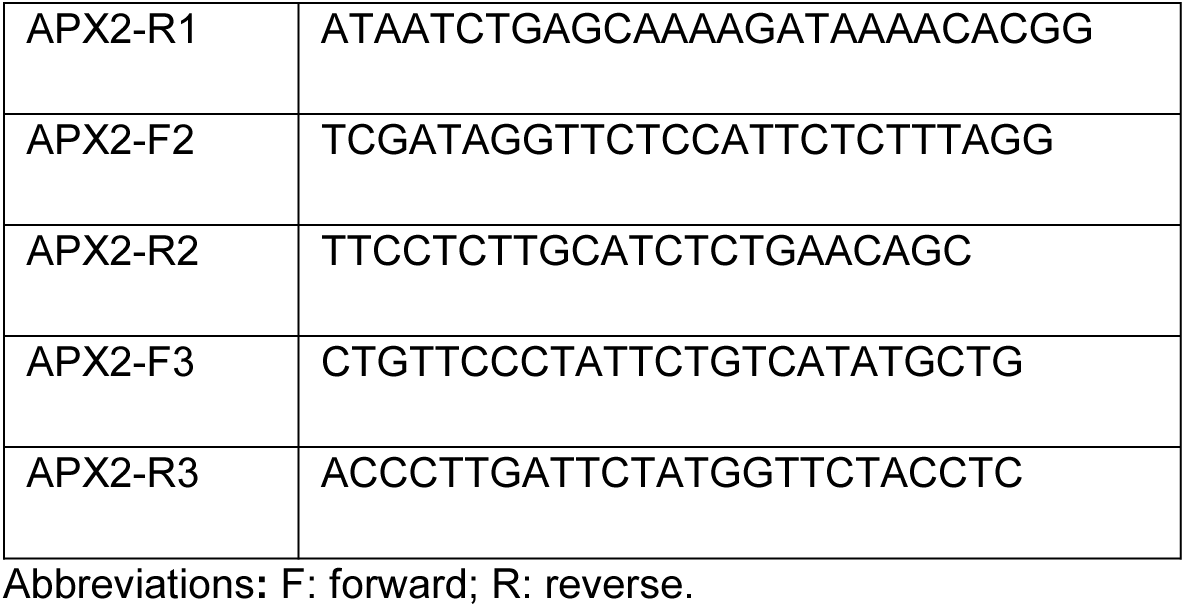
Primers Used For Chip-qPCR.

## Results

### Generation of *usp1* mutant lines

Two independent *Arabidopsis* knockout mutant lines, *usp1-1* and *usp1-2,* were generated by the CRISPR-Cas9 system utilizing two sgRNA targeted at *USP1* to study the protein function. The mutation/indels were determined by sequencing, which showed an addition of adenine in the mutant alleles of various lines (Fig. 1A). No morphological differences between the mutants and the wild-type plants were observed (Fig. 1B).

**Figure 1.**
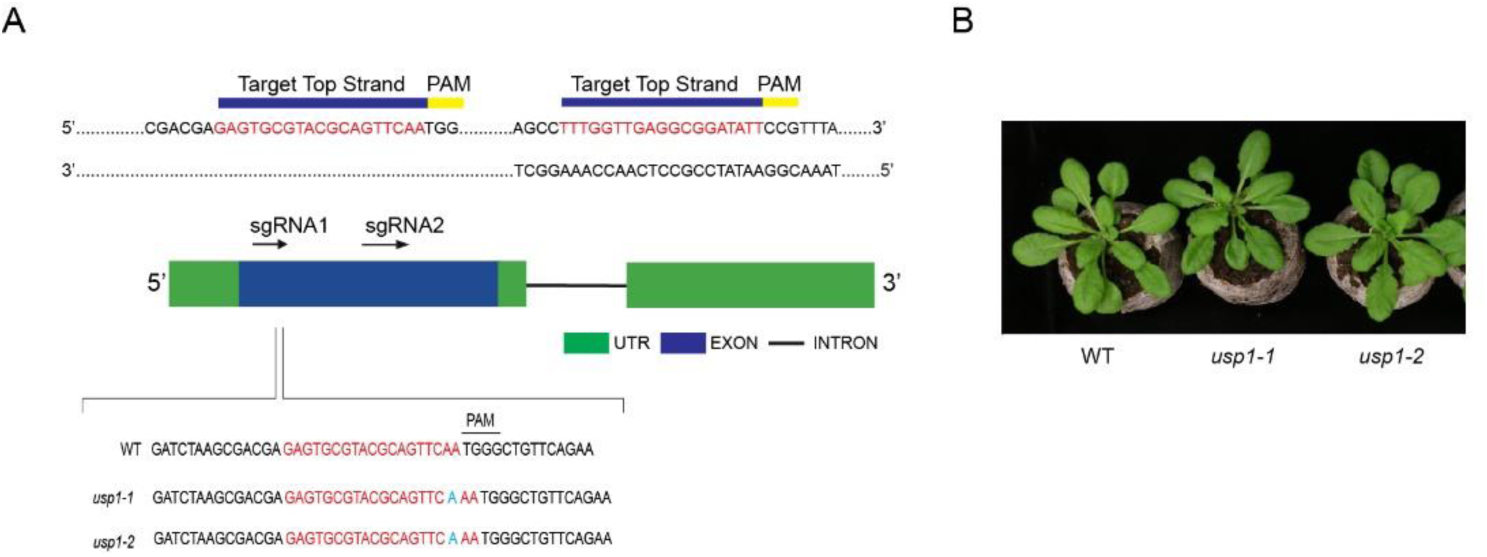
Characterization of *usp1* mutants. (**A**). CRISPR-Cas9 genome-editing strategy in *Arabidopsis* with two sgRNA targeting *USP1* and sequencing reads from selected T0 transformants showing the addition of adenine in various mutant alleles. (**B**). The morphological phenotype of WT (Col-0), *usp1-1*, and *usp1-* 2 mutants is identical for plants grown in jiffy peat pellets for four weeks.

### *usp1* mutants showed reduced acquired thermotolerance

To investigate the role of *USP1* in heat stress, we tested three sets of plants under different temperature regimes: untreated NHS (22°C), primed (37°+44°C), and non-primed acute heat stress (44°C) plants. For acute heat stress, plants were grown for 11 days before challenging them at 44°C for 30 min. Thereafter, plants were incubated at 22°C for five days before phenotypic analysis (Fig. 2A). Under these conditions, *usp1* mutant plants did not behave differently from WT plants (Fig. 2B). For acclimation, plants were primed at 37°C for 3 h and then allowed to recover at 22°C for two days followed by an acute heat stress at 44°C for 30 min and subsequent incubation at 22°C before phenotypic analysis. Consistent with previous findings, our phenotypic analysis of WT Col-0 plants demonstrated that 37°C primed plants exhibited significantly higher survival rates and fresh weight compared to non-primed plants to an acute heat stress at 44°C, indicating the beneficial effect of heat priming on thermotolerance. However, this priming effect was notably absent in the *usp1* mutant plants. The *usp1* mutants showed susceptibility to heat stress regardless of whether they had undergone priming, displaying comparable levels of damage and reduced growth under both temperature regimes. This suggests that *USP1* plays a critical role in acquired but not basal thermotolerance (Fig. 2B and C). Since *HSFA2* is the key regulator of heat stress memory, we also tested *hsfa2* mutant plants. Although *hsfa2* plants responded normally to acute heat stress, these plants were completely compromised to respond to the 37°C priming treatment (Fig. 2B and C). Importantly, *usp1* mutants behaved identical to *hsfa2* mutants both with respect to an acute heat stress and with respect to priming (Fig. 2B and C), indicating that *USP1* functions on the same pathway as *HSFA2*. The fresh weight of the *usp1-1* mutants was clearly compromised in 37°C priming (compared with control plants, with a decrease of 42.86% (Fig. 2D). The data were reproducible among three experiments, which were performed at different times. These results suggest that *USP1* is involved in the priming mechanism of acquired thermotolerance but not in the acute heat shock response in *Arabidopsis*.

**Figure 2.**
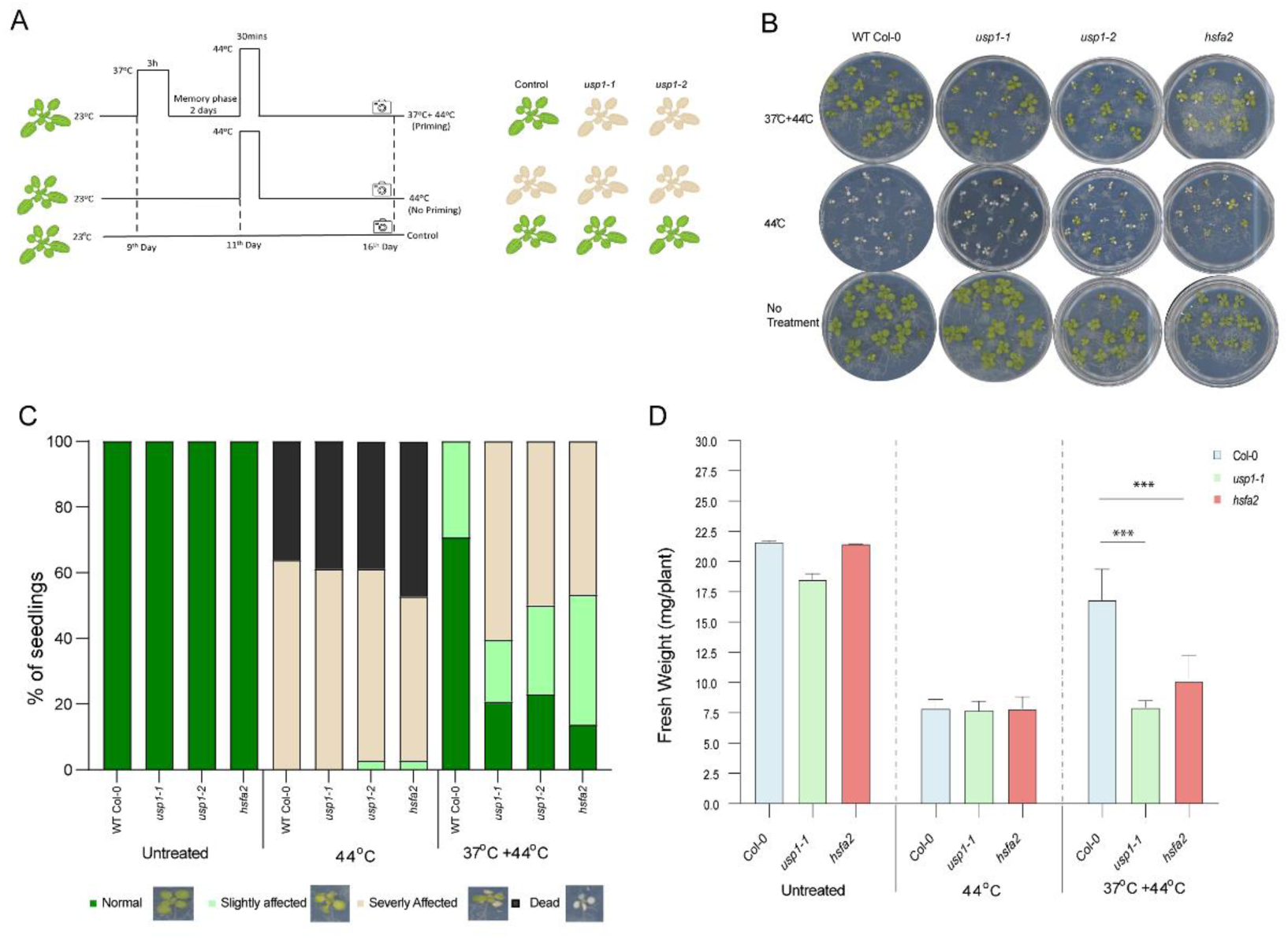
Physiological response of *usp1-1* mutants to basal and acquired thermotolerance. (**A**). Treatment scheme for heat stress memory: Top - priming: 9 day old plants were exposed to priming HS (37°C for 3 hours) then transferred to 22°C for 2 days before acute heat stress ( 44°C for 30 min) and recovery at 22°C for 5 days. Middle - no priming: 11 day old plants were treated at 44°C for 30 mins and incubated for recovery at 22°C for 5 days, Bottom - control plants were grown in parallel at 22°C for 16 days. Survival was scored after 5 days of acute heat stress treatment. (**B**). Representative images of heat stress memory assays (**C**). Distribution of phenotypic categories observed in the heat stress memory assay shown in B. (E). Fresh weight of Col-0 and *usp-1-1* mutant (16 days). The values under fresh weight represent the mean of at least twelve individual plants of each line. The error bars represent SE (standard error); one asterisk (*) indicates p < 0.05, two asterisks (**) indicate p < 0.01, three asterisks (***) indicate p > 0.001, ns indicates not significant as per Student’s t-test.

### Transcriptome Analysis of *usp1* mutants

To explore the regulatory mechanisms underlying *USP1*-mediated heat stress tolerance, a comparative transcriptome analysis of *usp1-1* mutant and WT plants was performed. Samples from three independent biological experiments were collected for RNA extraction and RNA-seq was carried out on an Illumina HiSeq instrument. A total of 428 differentially expressed genes (DEGs) were identified in WT and *usp1-1 mutant* plants with a two-fold change, p-value ≤ 0.05, q-value ≤ 0.05. Among these genes, 398 (93%) were upregulated, whereas 30 genes (7%) were downregulated (Fig. 3A). The upregulated genes were mainly enriched in GO terms of protein folding, electron transport, ATP biosynthetic, photosynthesis, and oxidative phosphorylation. No GO terms were identified in the downregulated genes. However, two downregulated genes in the *usp1-1* mutant (AT1G78370 and AT3G09270) encode glutathione transferases, which are implicated in combating ROS generation. Moreover, several downregulated CYP genes (AT2G42250, AT4G15393, AT1G13710, AT3G52970, and AT5G06905) are involved in protecting plants from abiotic stresses such as heat.

**Figure 3.**
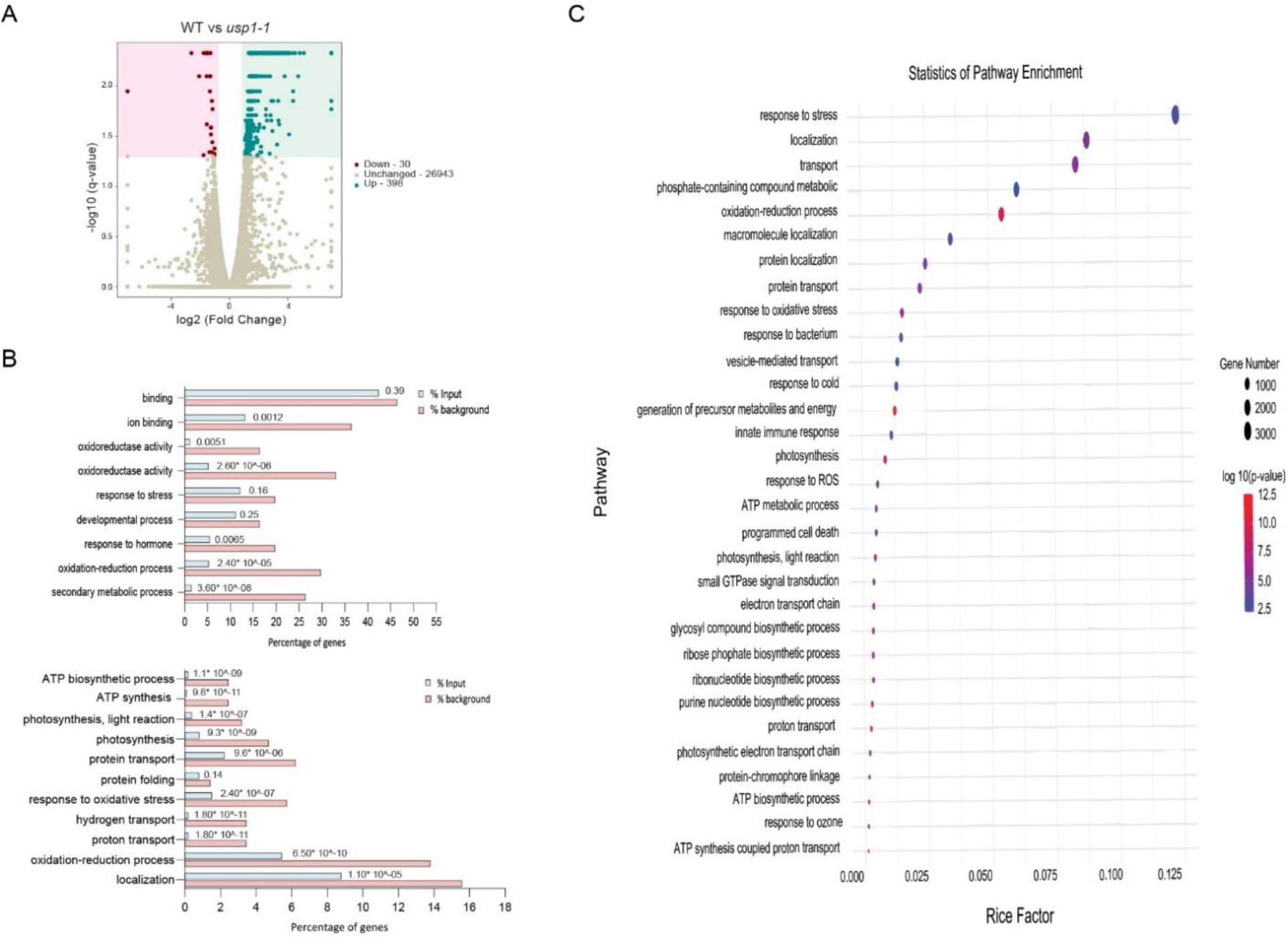
Transcriptome analysis of *usp1-1* mutant. (**A**). Volcano plots of different comparisons among WT mock and *usp1* mock. Log_2_ fold change is plotted against log_10_ (P-value). (**B**). Characteristics of differentially expressed genes (DEGs) in *Arabidopsis usp1-1.* Histograms represent functional enrichment of DEGs between Arabidopsis Col-0 (WT) mock and *usp1-1* mock in the GO databases. Top, Downregulated DEGs; Bottom, Upregulated DEGs. Blue represents input, and orange represents the reference control. The fold enrichment was determined based on the frequency of the genes associated with the term in comparison to their frequency in the genome. (**C**). KEGG enrichment analysis for the upregulated DEGs. R package ggplot2 was used to visualize the data. The size of the dot represents the number of DEGs, and the color indicates the corrected with a p-value of 0.05 of the pathway.

Most of the enriched genes from the upregulated cluster of genes showed clear enrichment of genes and strong associations with key metabolic pathways and cellular processes. Specifically, genes related to membrane protein folding (including members of the DNAJ superfamily, *ndhA*, *nad4*, and *nad6*), electron transport (e.g., *ndhG*, *ndhH*, *ndhJ*), ATP biosynthesis (*atpA*, *atpF*, and ATP synthase), photosynthesis (*psbH*, *psbC*, *ycf4*, *psbA*, and *psbE*), and oxidative phosphorylation (*atp1*, *atp6*, *atp8*, *ndhA*, *C*, *G*, and *ATP3*) were prominently enriched (Fig. 3B and C). The major genes are connected to energy metabolism linked to chloroplasts and mitochondria, suggesting that *USP1* plays an essential role in the ATP and NADPH metabolism. The genes in these pathways are crucial for cellular energy homeostasis under stress conditions where inter-organelle communication is essential. ATP involved in biosynthetic and oxidative phosphorylation pathways has been shown to respond dynamically to changes in the photosynthetic efficiency and redox balance in the chloroplast, influencing ATP production and ROS scavenging in the mitochondria. The electron transport components, especially *NADH* genes, support electron flow continuity, which is essential for balancing ATP/NADPH ratios in photosynthesis and adjusting energy transfer between chloroplasts and mitochondria to regulate energy levels (Yamori et al., 2016; Tikkanen et al., 2014; Shikanai et al., 2014).

Furthermore, DNAJ superfamily members function as molecular chaperones in protein folding and stabilization within chloroplast membranes, influencing photosynthetic efficiency. This intricate regulation of energy metabolism within the chloroplast and mitochondria maintains the cellular energy levels (Nishizawa-Yokoi et al.,2010; Lee et al., 2000). Overall, the differential expression of these genes may explain the role of *USP1* in abiotic stress, such as heat and oxidative stress.

### *USP1* regulates reactive oxygen species (ROS) levels during heat stress

Given the different roles of *USPs* in heat and oxidative stress (Jung et al., 2015; Chi et al., 2019 and Wu et al., 2023), we further investigated the role of *USP1* under 44°C heat stress (HS) by RT-qPCR analysis. The expression of *USP1* was significantly increased in heat-stressed WT plants compared to the non-heat stressed (NHS) plants (Fig. 4A). To analyze the accumulation of ROS in mutant and control plants, we examined ROS accumulation by 3,3-diaminobenzidine (DAB) and nitro blue tetrazolium (NBT) staining to detect H_2_O_2_ and O_2_^−^, respectively. DAB staining showed higher accumulation of hydrogen peroxide levels in the *usp1* mutant compared to WT Arabidopsis plants. Similar to DAB, NBT staining revealed that superoxide levels are higher in *usp1* mutant than in WT plants (Fig. 4B and C). Together, these results suggest that both superoxide and hydrogen peroxide levels are regulated by the *USP1*.

**Figure 4.**
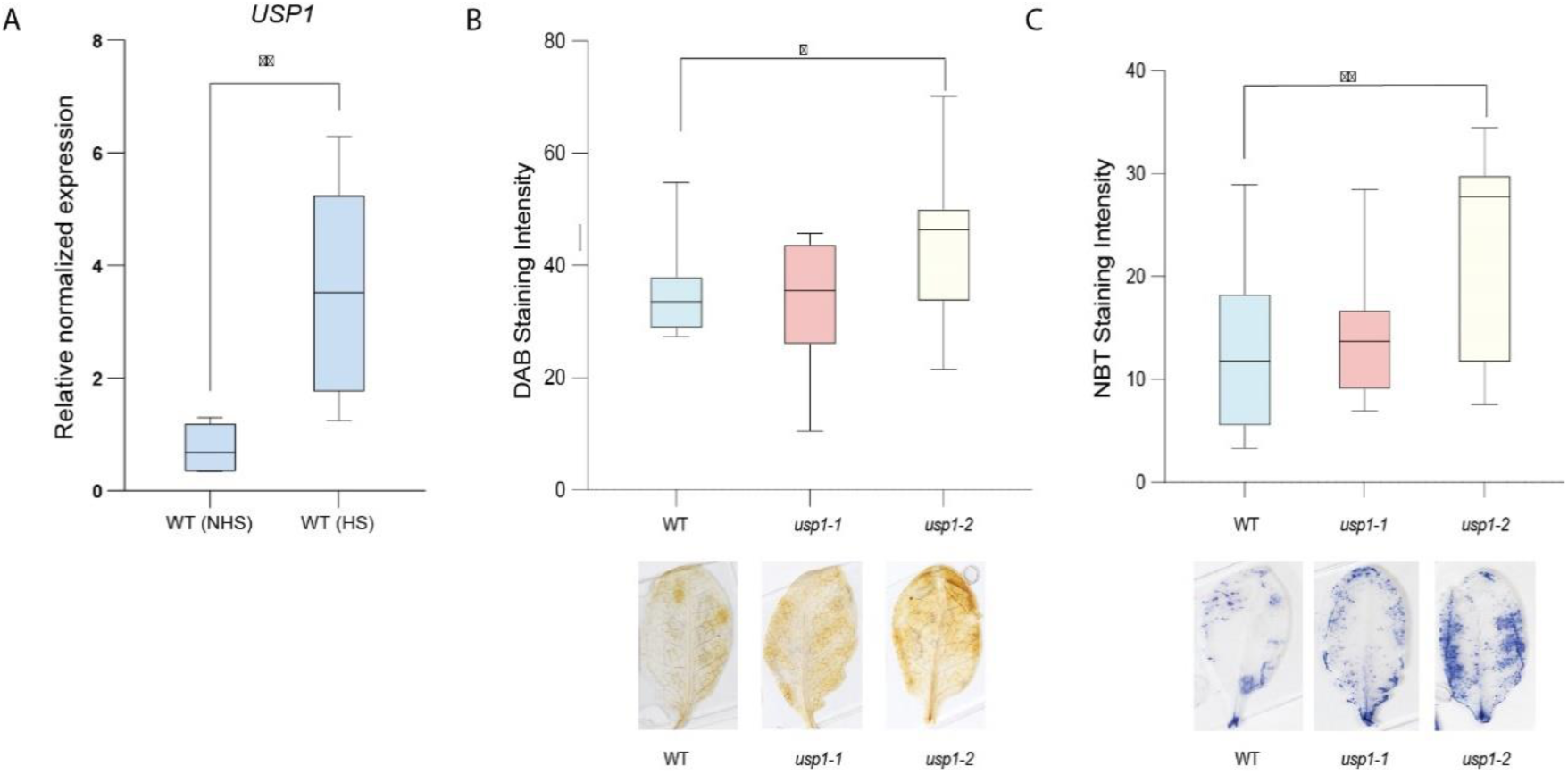
Effect of heat stress on USP1 and ROS levels in WT and *usp1* mutant plants. (**A**). Expression analysis of *USP1* in Arabidopsis WT plants non heat stress (NHS) and heat stress (HS) treatment via RT-qPCR. 11 day old plants grown at 22°C were treated at 44°C for 30 mins before recovery at 22°C for 1h before sample collection. Signal intensity for *USP1* transcripts was normalized with actin and ubiquitin. The expression levels of actin and UBQ10 were used for normalization. The error bars represent standard error, one asterisk (*) indicates P<0.10, (**) P<0.05, (***) P>0.001, Student’s *t-test*. The y-axis shows the difference in levels of gene expression plotted using a log2 scale. Two independent experiments were used (n=18). (**B and C**). Evaluation of H_2_0_2_ levels by 3,3’- diaminobenzidine staining (DAB) in WT and *usp1* mutant plants and O_2-_ levels by Nitro Blue Tetrazolium (NBT) staining. Three independent experiments (n=9, for each experiment). Data represent the error bars (SE, standard error, * indicates P<0.05, *** P>0.001) as per Student’s *t*-test.

### USP1 exhibits molecular chaperone activity on HSFA2

In previous studies, USPs were suggested to play a potential role as chaperones (Chuang et al., 2006 and Chi et al., 2013). To examine the role of USP1 as a chaperone, we tested the aggregation level of Malate Dehydrogenase (MDH) as a model ligand for USP1 at 25°C and 45°C. The USP1 protein reduced the heat-induced aggregation of MDH at 45°C compared to the MDH alone (Fig. S4). Since the genetic analysis indicated that USP1 functions on the same pathway as HSFA2, we investigated what biochemical function might underlie this process. For this purpose, we tested whether USP1 possesses molecular chaperone activity *in vitro*. Differential scanning fluorimetry (DSF) has been widely used to study the protein thermal stability, and we utilized SYPRO orange as a sensing dye with the direct use of qPCR (Malik et al., 2013 and Zbacnik et al., 2017). We performed DSF using LDH as a model ligand for USP1 (Fig. 5A). Using thermal destabilization, the DSF test has been widely utilized to deduce chaperone activity. In the absence of USP1, LDH was destabilized at 45°C. The addition of USP1 protein caused a positive shift of 0.5°C in the melting curve of HSFA2 (Tm, app = 58 + 0.5°C). In the absence of USP1 destabilization of HSFA2 was observed at 64°C (Fig. 5B). These data show that USP1 can stabilize HSFA2 by binding to the native folded HSFA2 protein. The difference in heat-primed WT and *usp1-1* plants was recapitulated by the fact that it did create a substantial thermal shift of HSFA2 (Fig. 5B). To analyze the protein-protein interaction of USP1 and HSFA2, we performed *in silico* protein-protein docking using pydock. The best docking poses are visualized using PYMOL (Fig. S5). Fig. S5 is a PYMOL output that shows all hydrogen-bonding interactions contribute to HSFA2 association. We then used a computational method to calculate the binding energy between USP1 and HSFA2 using Vander Waal potentials, electrostatic potential, desolvation energy, and total binding energy. The total binding energy of the interaction is low, as shown in table 5, indicating more critical residue interactions between both proteins (Zhang et al., 2020; Table 5).

**Figure 5.**
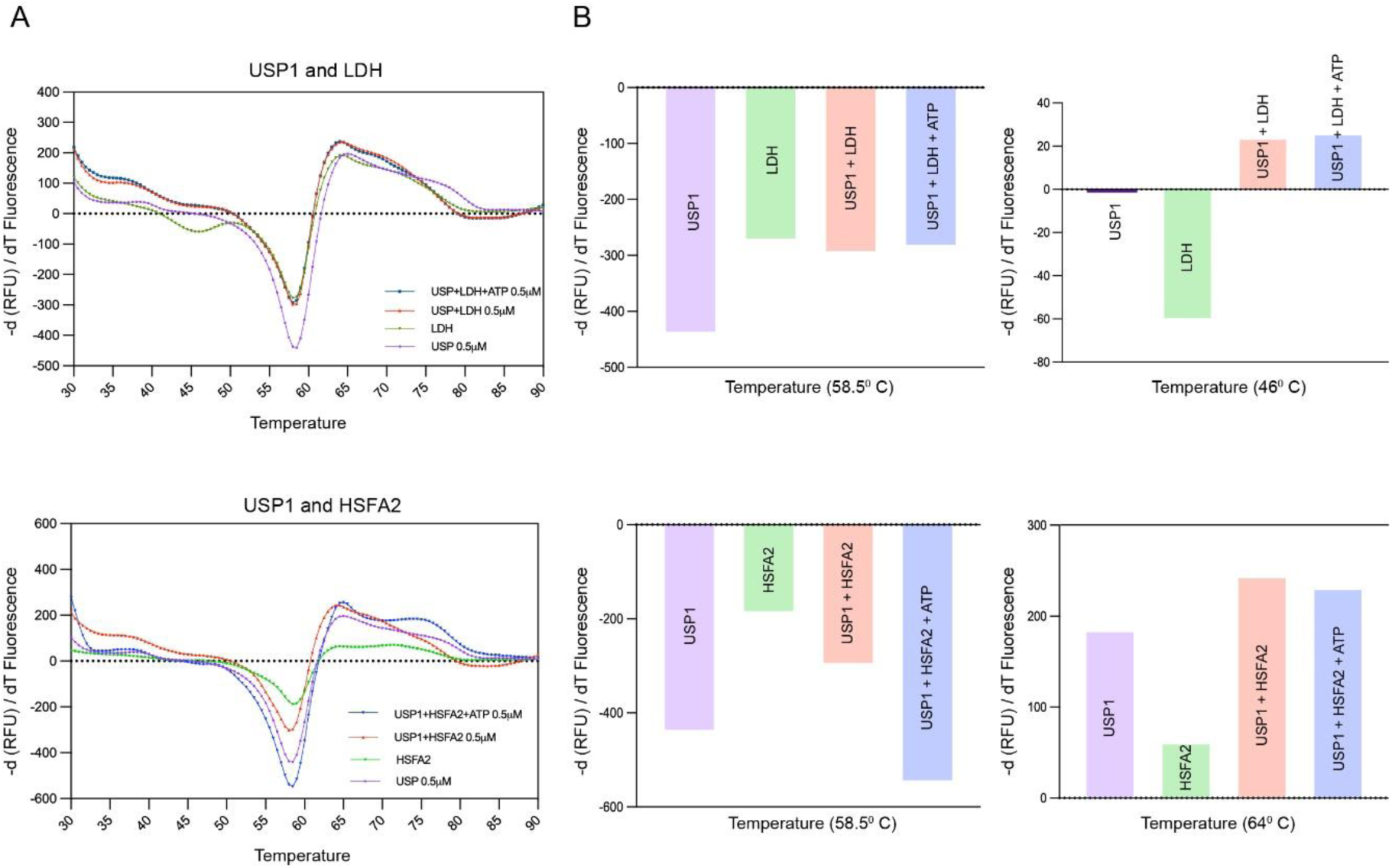
USP1 binding to HSFA2, as probed via DSF. (**A**). DSF melting curve shows protein unfolding by increase temperature, accompanied by an increase in SYPRO fluorescence. The melting temperature (Tm) values were determined by calculating the first derivative from the melting curve with the Biorad CFX manager software. Data are represented in negative derivative of RFU with respect to temperature. The protein target is USP1, positive substrate LDH, ligand HSFA2, where present, are 0.5 µM concentration. The total reaction volume of each well is 10 µL. (**B**). The X-axis value temperature that corresponds to the Y-axis value (-d(RFU)/dT) in the resulting curve resembles Tm. All data were collected in triplicate.

**Table 5.** The tabular column depicts the Top 3 ranks (pydock and HDOCK)

| **Complex** | **Electrostatic energy (kcal/mol)** | **Desolvation energy (kcal/mol)** | **VdW (kcal/mol)** | **Total Binding energy** |
| --- | --- | --- | --- | --- |
| USP1 and HSFA2 | -5.320 | -32.533 | 45.585 | -33.295 |
|  | -11.819 | -22.736 | 20.615 | -32.493 |
|  | -13.228 | -22.182 | 31.950 | -32.215 |

| **Complex** | **Docking Score** | **Confidence score** | **Ligand rmsd (Å)** |
| --- | --- | --- | --- |
| USP1 and HSFA2 | -227.54 | 0.8250 | 34.19 |
|  | -225.21 | 0.8182 | 35.11 |
|  | -217.16 | 0.7930 | 32.03 |

Abbreviations: rmsd- root mean square deviation; VdW- Vanderwaals

### USP1 regulates the expression of heat stress signature genes

We investigated the role of USP1 in regulating heat stress memory and heat-responsive genes by analyzing transcript levels. qRT-PCR was performed on heat-responsive and memory genes in *usp1-1* mutants and WT plants after a 30-minute exposure to 44°C, comparing primed (37°C+44°C), and direct heat stress conditions (44°C) (Fig. 6A). The results show that in WT the transcript levels of *HSFA2*, *HSFA6B*, *HSFA8*, *HSFA7a*, and *HSFA3* were significantly higher in primed compared to non-primed plants; in contrast in *usp1-1* mutant plants, the expression of all the above listed HSFs did not change significantly under primed compared to non-primed conditions (37°C+44°C and 44°C) (Fig. 6B). Similarly, *USP1* is crucial for the induction of heat stress memory genes like *APX2* and *HSP18.2*, as well as heat stress response genes such as *BIP3*, *HSP90.1*, *HSP70*, and *HSP101* (Fig. 6B) as the expression of all the genes did not change for *usp1-1* mutant plants like WT plants in primed compared to non-primed conditions.

**Figure 6.**
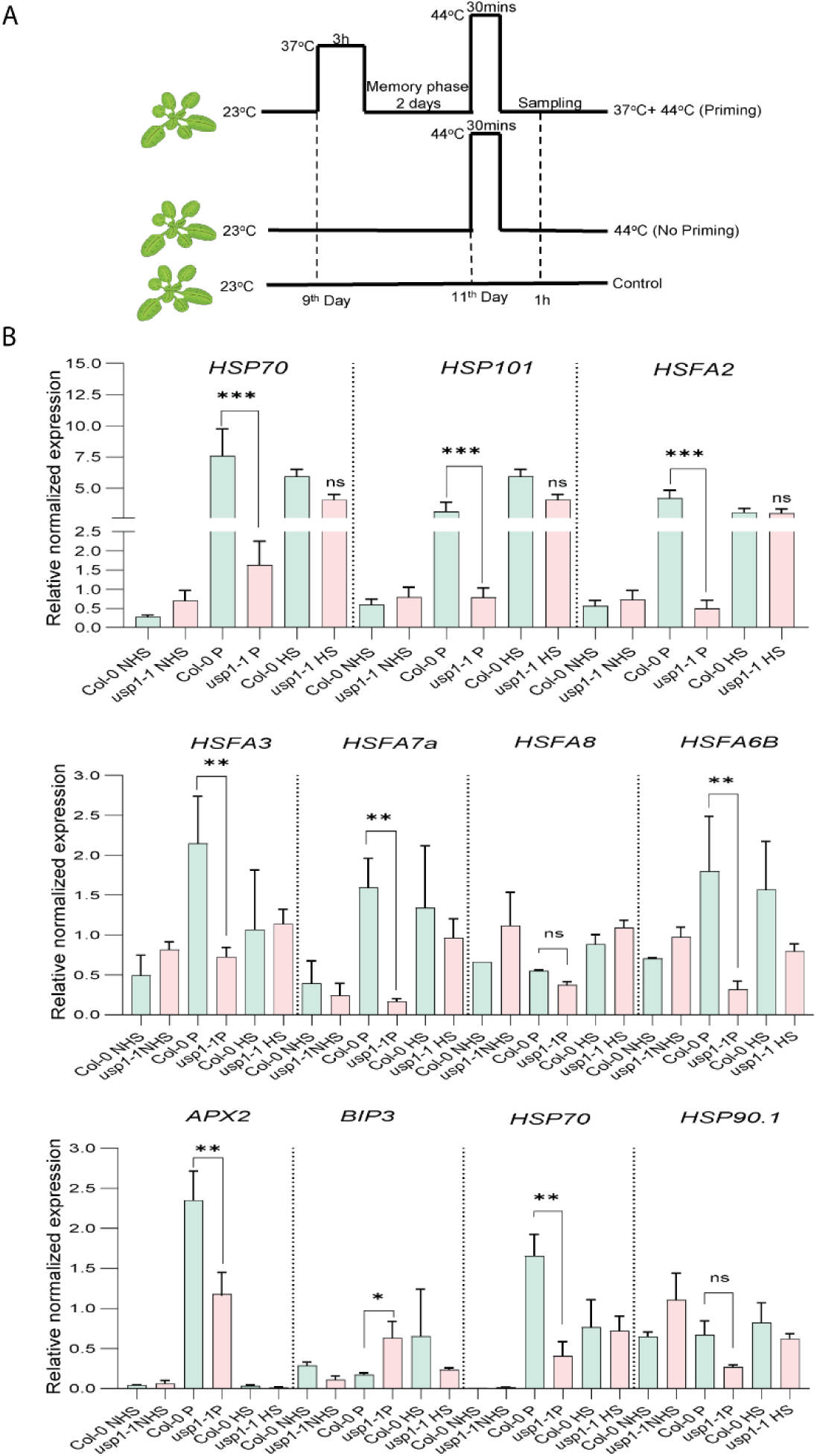
Thermotolerance is associated with higher expression of heat-responsive genes. (**A**). Treatment scheme for heat stress memory: plants are exposed to either priming HS (37°C for 3 hours), then after a recovery period of 2 d at 22^0^C, plants received an acute heat treatment for 30 min (44°C) before a second recovery at 22^0^C. As a control, plants were kept at 22^0^C, and non-primed plants were grown for 11 days at 22°C before treatment at 44°C for 30 mins and subsequent recovery at 22°C.Transcript levels of *HSFA2*, *HSP101* and *HSP18.2.* (**B**). Transcript levels of heat shock factors (*HSP70, HSP101, HSFA2, HSFA3, HSFA7a, HSFA8, HSFA6B, APX2, BIP3, HSP70 and HSP90.1*) were measured by qRT-PCR in control, 37^0^C heat-primed, 37^0^C heat-primed and 44^0^C heat-stressed, and 44^0^C heat-stressed at 1 h of recovery at 22^0^C in WT and *usp1* mutant plants. Vertical bars indicate the standard error for three biological replicates with 16 plants per treatment. The error bars represent SE (standard error); one asterisk (*) indicates p < 0.05, two asterisks (**) indicate p < 0.01, three asterisks (***) indicate p > 0.001, ns indicate not significant as per Student’s t-test.

These findings indicate that *USP1* plays a key role in regulating the expression of heat stress signature genes for acquired thermotolerance. Since HSFA2 is known to be essential for thermomemory, we further explored the interaction between USP1 and HSFA2 proteins through docking analysis and visualization using PYMOL (Fig. S5). This analysis corroborates the gene expression patterns observed, highlighting a compromised heat stress response in thermoprimed and heat-stressed *usp1-1 plants*.

### USP1 boosts H3K4me3 thermopriming levels at the *APX2* and *HSP18.2* gene loci

To investigate the role of USP1 in heat stress memory, we performed chromatin immunoprecipitation followed by quantitative PCR (ChIP-qPCR) to measure the levels of H3K4me3, a histone mark associated with active transcription, in primed (P) and non-heat stress (NHS) plants. The priming treatment involved subjecting plants to 37°C for 3 hours, followed by a recovery period of 24 hours at 22°C (Fig. 7A). In line with previous findings (Lämke et al., 2016; Liu et al., 2018), thermoprimed WT plants exhibited a significant enrichment of H3K4me3 at specific regions of the heat stress-responsive *APX2* and *HSP18.2* genes, specifically at regions 2 and 3 of *APX2* and region 2 of *HSP18.2* (Fig. 7B & C). This enrichment is indicative of the establishment of a transcriptionally active chromatin state that supports the rapid reactivation of these genes upon subsequent heat stress exposure.

**Figure 7:**
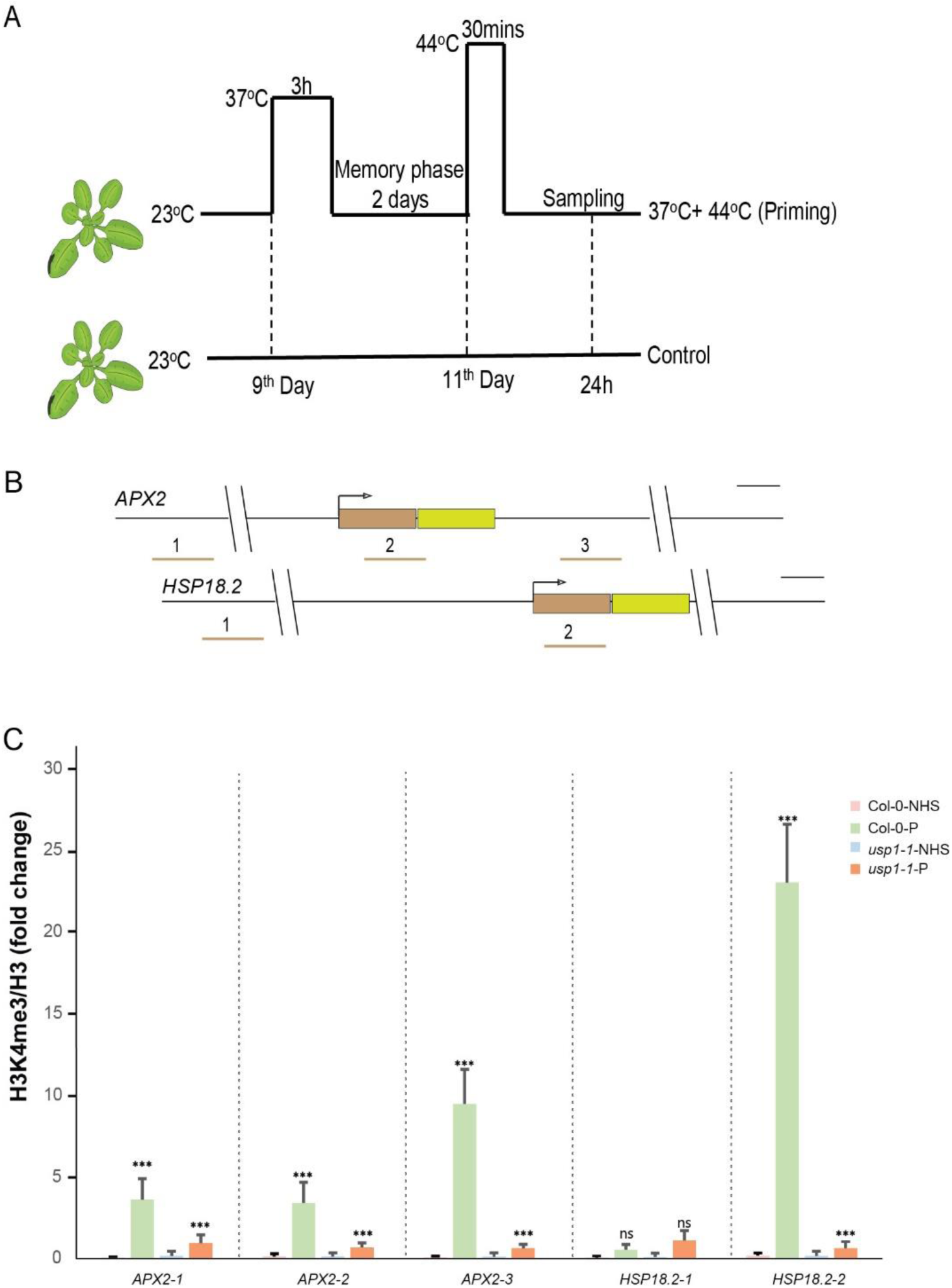
USP1 is required for heat stress memory and acquired thermotolerance. (**A**). Treatment scheme for heat stress memory: plants are exposed to priming HS (37°C for 3 hours) 2 d after (44°C for 30 min). Middle, plants without priming: 11-day-old plants that were grown at 22°C were treated at 44°C for 30 mins and incubated for recovery at 22°C. Bottom, control plants were grown in parallel at 22°C for 11 days. (**B**). *APX2* and *HSP18.2* gene models drawn to scale (brown, 5′ untranslated region; light green boxes, exons; angled arrow, transcription start site). The underneath numbers with gray bar indicate the positioning of regions analyzed for ChIP-qPCR, three regions of *APX2* and two regions of *HSP18.2*. (**C**). Relative enrichment of H3K4me3 at *APX2* and *HSP18.2* in control non-primed (NHS) and 37°C-primed plants (P) at 24 after priming as determined by chromatin immunoprecipitation-qPCR for the indicated regions of *APX2* and *HSP18.2*. Amplification values were normalized to input and H3. The error bars represent SE (standard error); one asterisk (*) indicates p < 0.05, two asterisks (**) indicate p < 0.01, three asterisks (***) indicate p > 0.001, ns indicate not significant as per Student’s t-test.

In contrast, the *usp1-1* mutant plants did not exhibit a significant increase in H3K4me3 levels at these loci following the priming treatment (Fig. 7B & C). This lack of H3K4me3 enrichment suggests that USP1 is important for the epigenetic modification of heat stress memory gene loci. Therefore, USP1 plays a critical role in the establishment of acquired thermotolerance.

## DISCUSSION

In this study, we present evidence that USP1 is involved in the regulation of acquired thermotolerance in *Arabidopsis*. We show that USP1 acts as a molecular chaperone for HSFA2 that is required for acquired thermotolerance and sustained induction of heat stress memory genes. USPs were previously reported to play a role in abiotic stress in tomato, cotton, and rice (Fan et al., 2024). Here, we report that USP1 positively regulates the heat stress acclimation response in *Arabidopsis*. Mutants of *usp1* showed no phenotypic changes under normal growth conditions. However, although USP1 knock-out mutant plants behave like wild-type plants to acute heat stress at 44^0^C, they are totally compromised in acquired thermotolerance (Fig. 2). The transcriptome analysis of *usp1* mutant plants revealed massive upregulation of genes related to electron transport, ATP synthesis protection against ROS. The accumulation of ROS causes cellular damage, inducing the activation of antioxidants and ROS scavenging genes to maintain cellular redox homeostasis and HS acclimation (Choudhury et al., 2017). In response to elevated ROS levels, the expression of HS response genes, such as *MYBs*, *NACs*, and *HSPs*, is triggered to mitigate ROS-induced damage and enhance stress tolerance (Jagtap et al., 2023). Interestingly, among these genes, 50 were chloroplast and 19 mitochondrial-encoded genes. These genes function as alternative oxidase, ATP synthase, NADH dehydrogenases, and ribosomal proteins. These results highlight the role of organelle-related pathways in HS responses. Future studies should focus on unraveling the molecular mechanisms underlying these heat stress-specific functions to better understand their contributions to cellular resilience.

USPs are found across all organisms and play key roles in responding to various environmental stresses (Kerk et al., 2003). Consistent with TAIR and Bar eFP Browser data, *USP1* expression is significantly induced under heat stress. Thermotolerance and the ability to adapt quickly to changing temperatures are crucial for sessile plants. However, how USPs play a role in plant thermotolerance remains poorly understood. HSPs are activated by heat shock factors (HSFs) and play critical roles in plant heat tolerance (Lee et al., 2007; AH El-Sappah et al., 2022). HSPs act as molecular chaperones to stabilize and refold proteins under stressful conditions. For example, HSP101 has been identified as a crucial factor in acquiring thermotolerance in various plant species (Gurley et al., 2000), and other studies have shown that proteins like OsHSP101 interact with OsHSP16.9A to confer thermotolerance in rice (Liu et al., 2023) and calcium-dependent protein kinase (ZmCDPK7) in maize mediating the phosphorylation of sHSP17.4 that regulate heat stress responses, including thermotolerance and heat memory (Lin et al., 2014; Zhao et al., 2021).

Our molecular analysis of both wild-type and *usp1* mutant lines revealed that *usp1* mutants can activate a typical HS response upon exposure to acute heat stress (44°C) but unlike Col-0, *usp1* plants are unable to sustain heat stress after acclimation at 37°C. This phenotypic response mirrors that of *hsfa2* mutants, which also fail to activate heat stress memory. Our findings highlight the critical role of USP1 as a molecular chaperone for HSFA2. Using differential scanning fluorimetry, we established that USP1 interacts directly with HSFA2, acting as a chaperone under thermal stress conditions (Fig. 5).

USP1 interaction with HSFA2 is important to maintain stability, prevent aggregation, and modulate its activation under high temperatures. Structural modeling further confirmed that USP1 occupies a binding pocket on HSFA2, highlighting the molecular basis of this interaction (Fig. S5). This interaction enables HSFA2 to remain functional in recruiting histone H3K4 trimethylation to activate gene expression (Charng et al., 2007; Fig. S6) thereby ensuring effective heat stress memory. These findings suggest that USP1 not only enhances HSFA2 stability but also supports its epigenetic regulatory roles. Recent research has highlighted the importance of chromatin modifications, particularly histone H3K4me3, in regulating heat-responsive memory genes such as *APX2* and *HSP18.2*, enabling plants to remember heat stress and respond more effectively to subsequent heat stress (Liu et al., 2018; Shekhawat et al., 2022). We observed that USP1 plays a crucial role in facilitating H3K4me3 enrichment at these loci, as *usp1* mutants failed to accumulate H3K4me3 at heat stress memory genes (Fig. 7). This points to USP1’s significant role in establishing and maintaining thermotolerance, likely through its interaction with HSFA2. Our findings also show that USP1 influences the expression of key heat response genes, such as *HSP90.1*, *HSP70*, and *HSP101* (Fig. 6 B& D). These proteins are involved in protein refolding and cellular protection, as well as in regulating reactive oxygen species (ROS) levels. The compromised expression of these genes in *usp1* mutants under primed treatment suggests that USP1 contributes to long-term adaptation to high temperatures. Interestingly, other members of the USP family in *Arabidopsis* have also been shown to exhibit chaperone functions, such as AtUSP, which protects intracellular molecules from heat and oxidative stress (Jung et al., 2015). *Atusp* mutants are compromised under acute heat stress, and overexpression of *AtUSP* enhances heat tolerance. AtUSP has also been reported to act as an RNA chaperone during cold stress, suggesting that USPs may have broader roles in stabilizing both protein and RNA molecules during temperature fluctuations (Kang et al., 2013; Melencion et al., 2017).

In summary, our study provides compelling evidence that USP1 plays a pivotal role in heat stress tolerance and memory in *Arabidopsis* by acting as a chaperone for HSFA2 and facilitating the accumulation of histone H3K4me3 at heat-responsive genes. In this context, the interplay of pre-and post-harvest thermal treatments emerges as promising strategies to mitigate post-harvest disorders such as superficial scald and to enhance fruit quality during storage by induction of HSFs and HSPs (Marc et al., 2020). Given the complexity and importance of USPs in managing protein and RNA stability during stress conditions, further research into this protein family could uncover novel mechanisms of plant stress resilience, with potential applications in developing crops with enhanced thermotolerance to adapt to climate change.

## Author Contributions

P.M., H.H., and N.R. designed the study. P.M., K.S., A.F., H.M. A, A.A.A., and N.R. performed experimental work. P.M. and N.R. performed *in silico* analysis and analyzed data. P.M., K.S., and N.R. wrote the paper. All authors have read and agreed to the published version of the manuscript.

## Data Availability Statement

RNA-Seq data are available at NCBI’s Gene Expression Omnibus GEO Series under accession number GSE286899.

## Acknowledgments

We would like to thank the KAUST Bioscience Core labs for technical assistance for RNA sequencing and all members of the Hirt lab.

---

**Figure S1.**
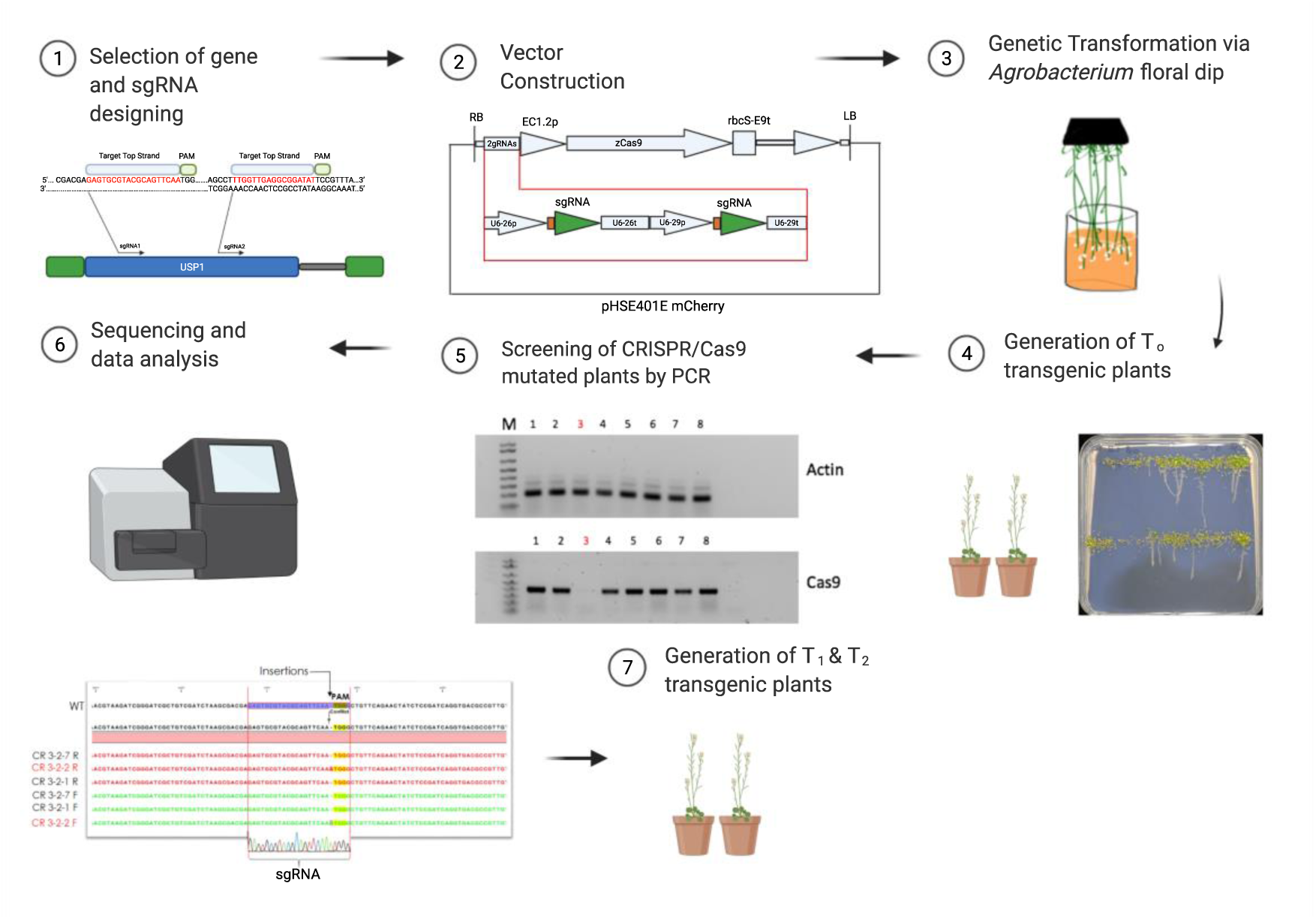
Workflow and Generation of CRISPR/Cas9 knockout mutant lines in Arabidopsis. Physical map of PHSE401 mcherry vector carrying two-gRNAs targeting two regions of Arabidopsis gene *AtUSP1.* The construct was transformed via Agrobacterium floral dip method. Screening of T1 transgenic plants on agar plates and select the survival to grow in the soil. The DNA of plants extracted, PCR amplified, and sequenced. Analysis of mutations using sequence alignment software to select for Indels and select homozygous mutants. Screening for T2 and T3 homozygous mutants and cas9 free lines is key steps to generate CRISPR/Cas9 lines.

**Figure S2.**
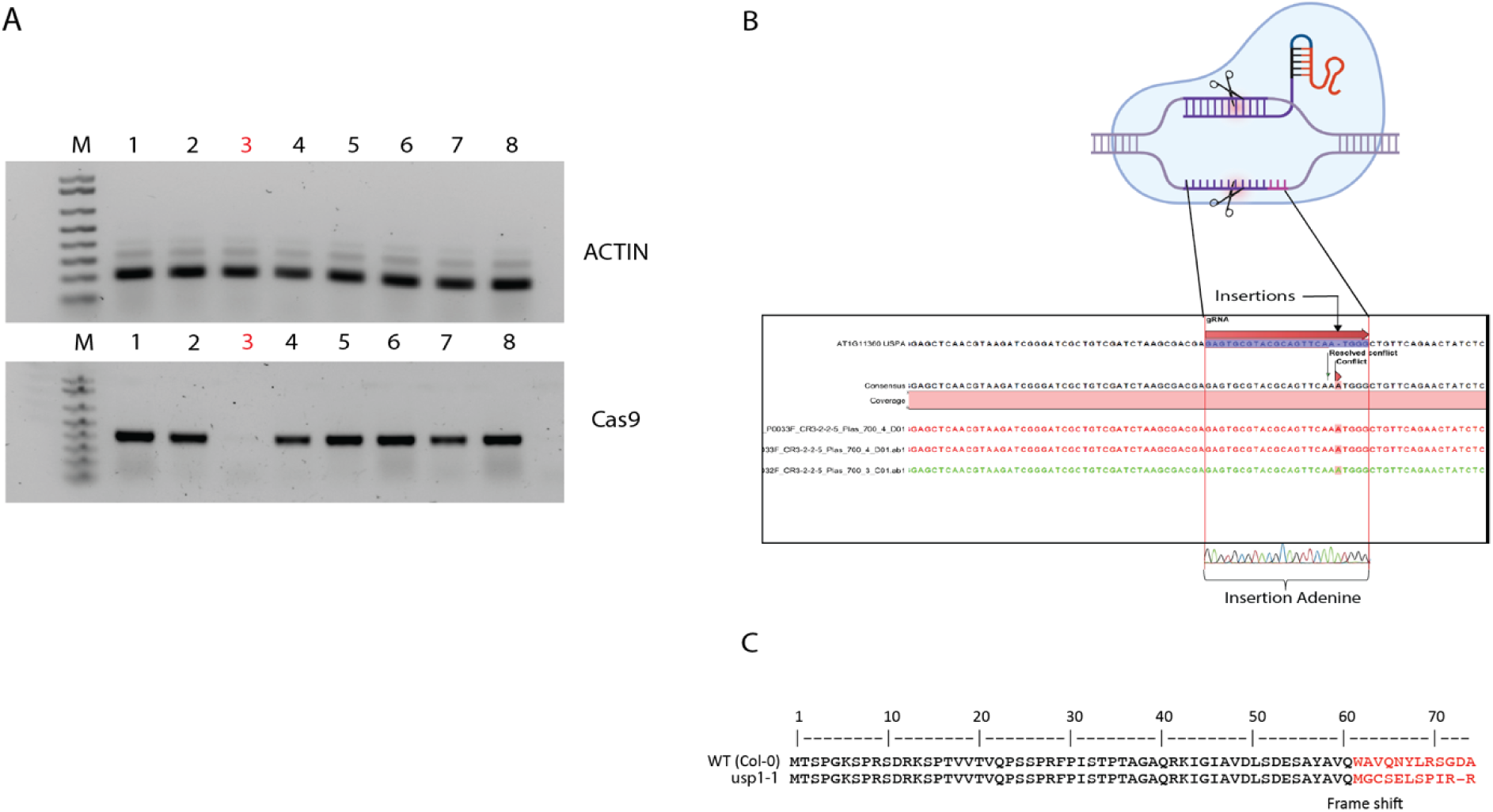
(**A**) Screening for Cas9 free CRISPR/Cas9 lines in subsequent generations. (**B**) *In silico* analysis of nucleotide sequence showing the frameshift indel resulting in a truncated protein. (**C**) One bp upstream of the PAM (TGA) within the sgRNA target sequence (highlighted), all causing frameshift and formation of premature stop codon.

**Figure S3.**
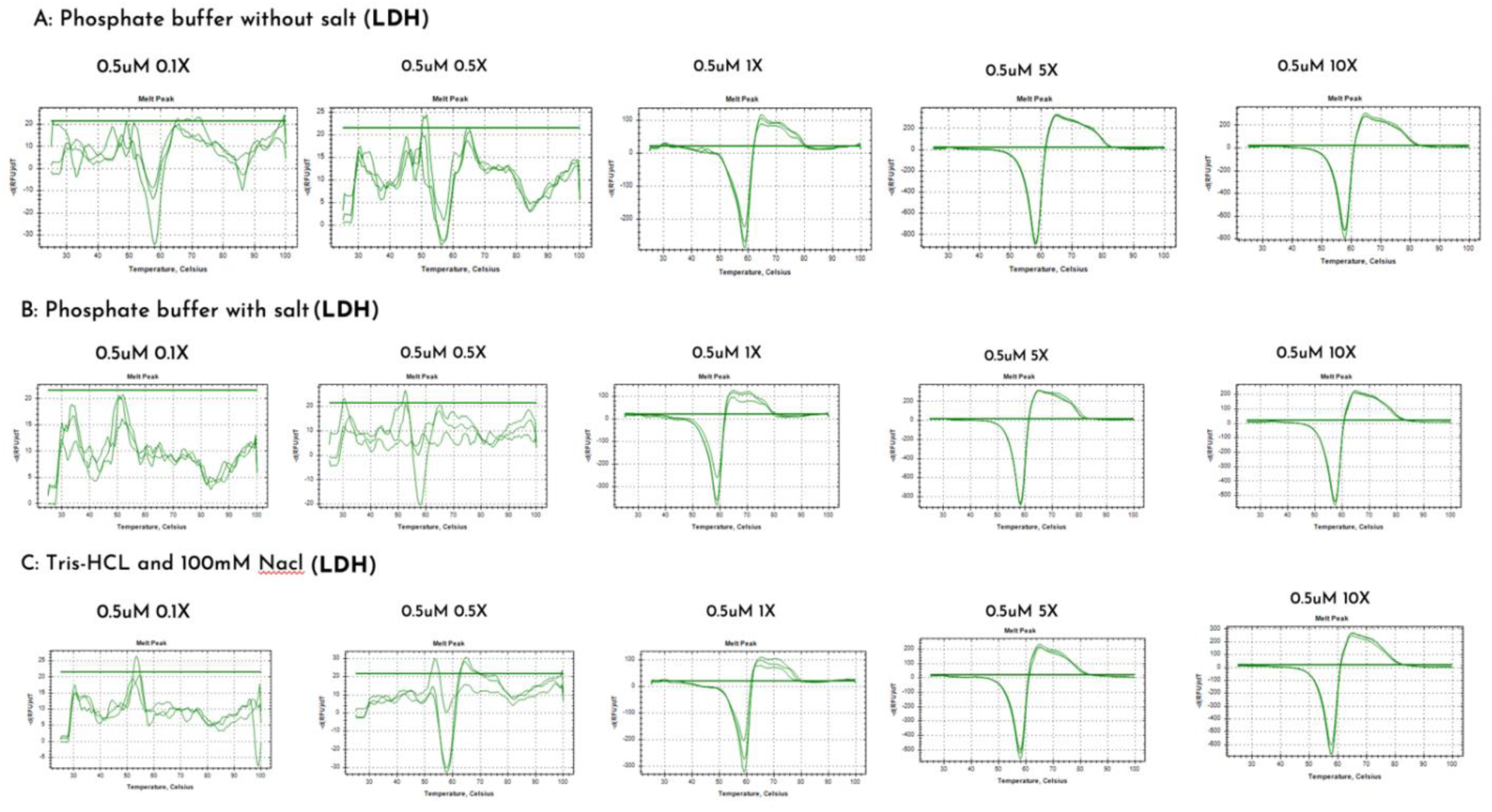
Differential scanning fluorimetry protein melting curves of the Lactate Dehydrogenase (LDH). Buffer standardization using DSF SYPRO orange fluorescence method indicates 0.5µM protein 1X, 5X, and 10X concentration of the different buffers is best suitable for the experiment.

**Figure S4.**
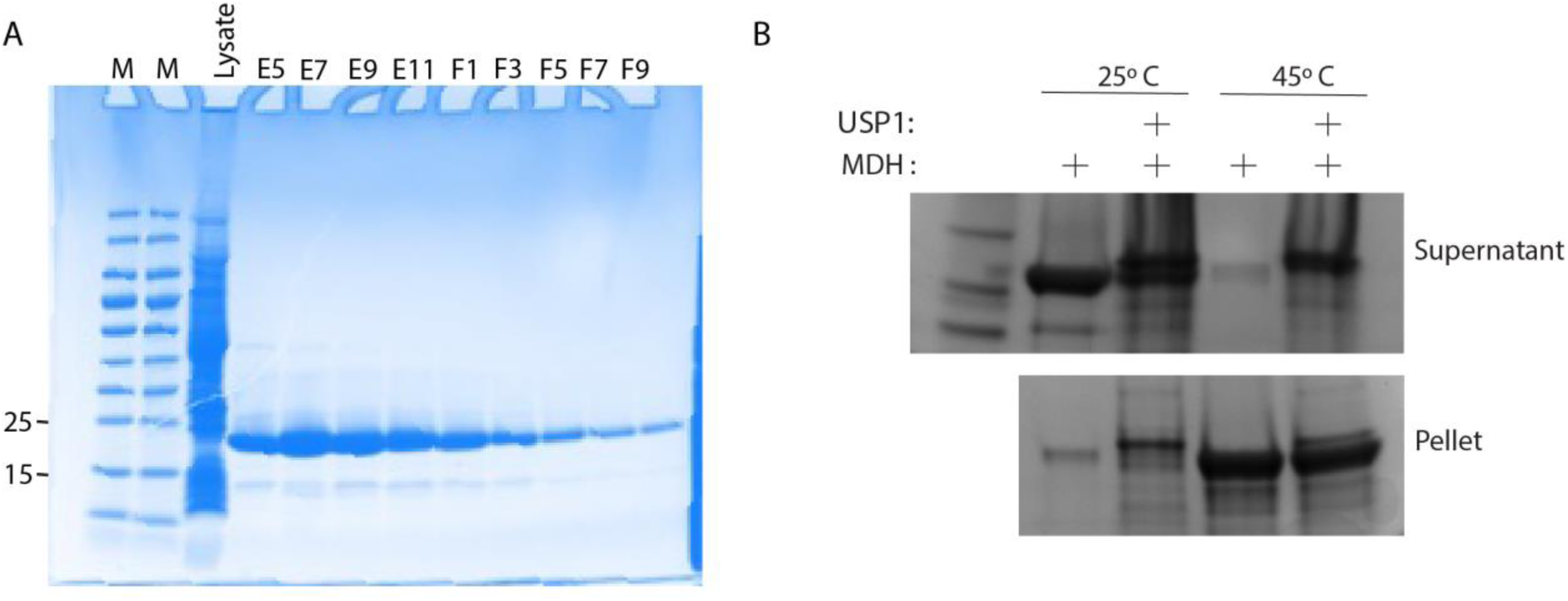
Chaperone activity of recombinant USP1 protein. (**A**) SDS-PAGE analysis of protein fractions from the AKTA purification of USP1. Lane 1 &2: Marker; lane 3: flow through after first wash; lanes 4-12, fractions from column after elution buffer was applied. USP1 protein (26.5 kDa). **(B**) The simplest type of activity test for a chaperone determines its capability to prevent the aggregation of a model substrate. Malate Dehydrogenase was incubated in the presence or absence of USP1 protein at 25°C and 45°C for 1 hour. After centrifugation, supernatant and pellet fraction were analyzed by SDS-PAGE.

**Figure S5.**
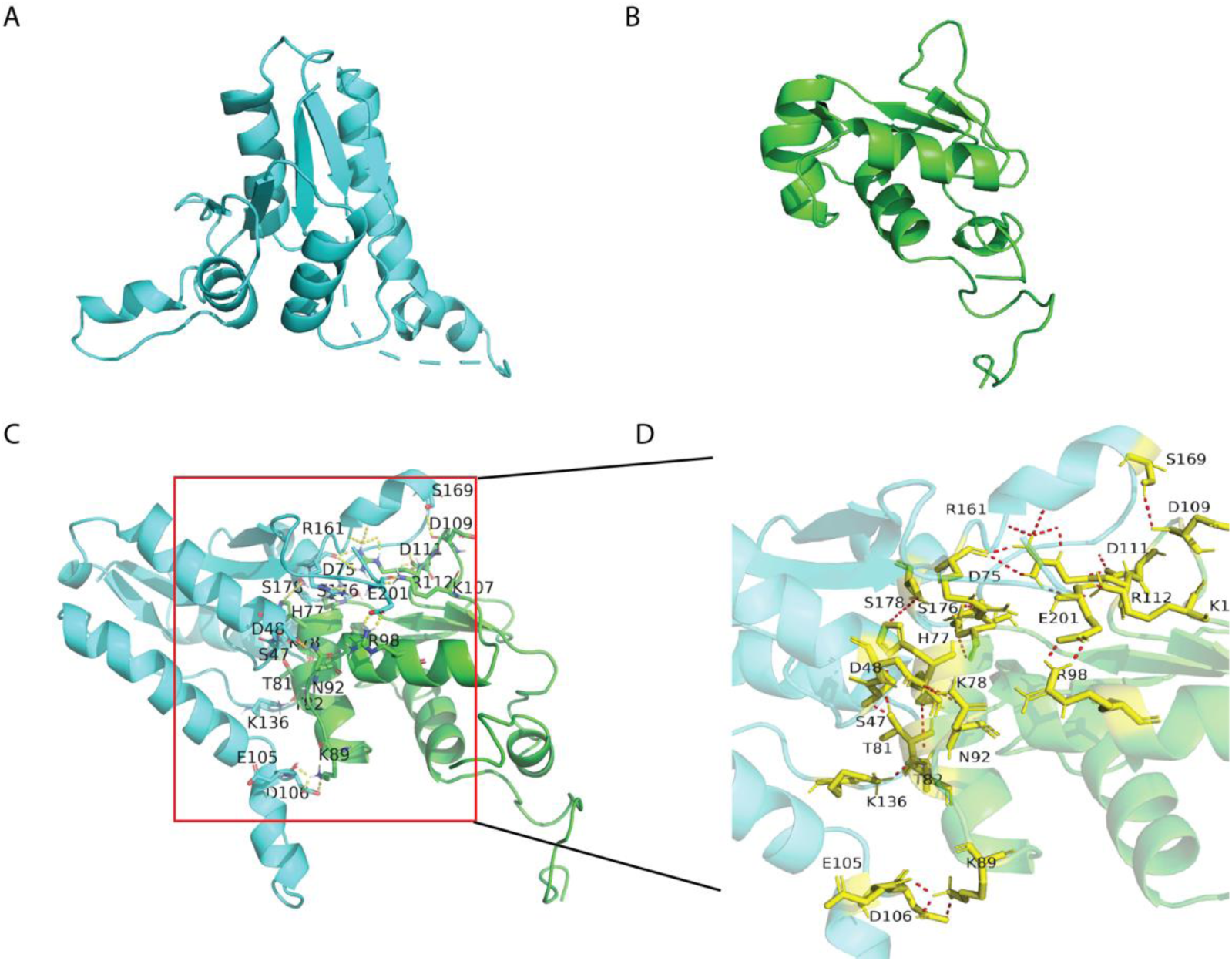
Structure and interaction of the USP1 and HSFA2. (**A**). Structure of the USP1 protein in light blue colour (Uniprot:PF00582; cyan), generated using Phyre2. (**B**). Crystal structure of HSFA2 (Uniport: HSFA2; green) in green colour. (**C & D**). Zoom into the HSFA2 binding pocket and USP1 interaction. Hydrophobic interaction: yellow sticks with red bonds; hydrogen bonds: dashed lines. For 3D structural analysis of the USP1 protein, Phyre2 (http://www.sbg.bio.ic.ac.uk/phyre2/) was used using the closest available crystal structure. Potential binding sites of USP1 and HSFA2 were analyzed by autodock vina using the generated Phyre2 3D structure with default parameters. Modelling and docking results were visualized using UCSF Chimera v1.12 software.

**Figure S6.**
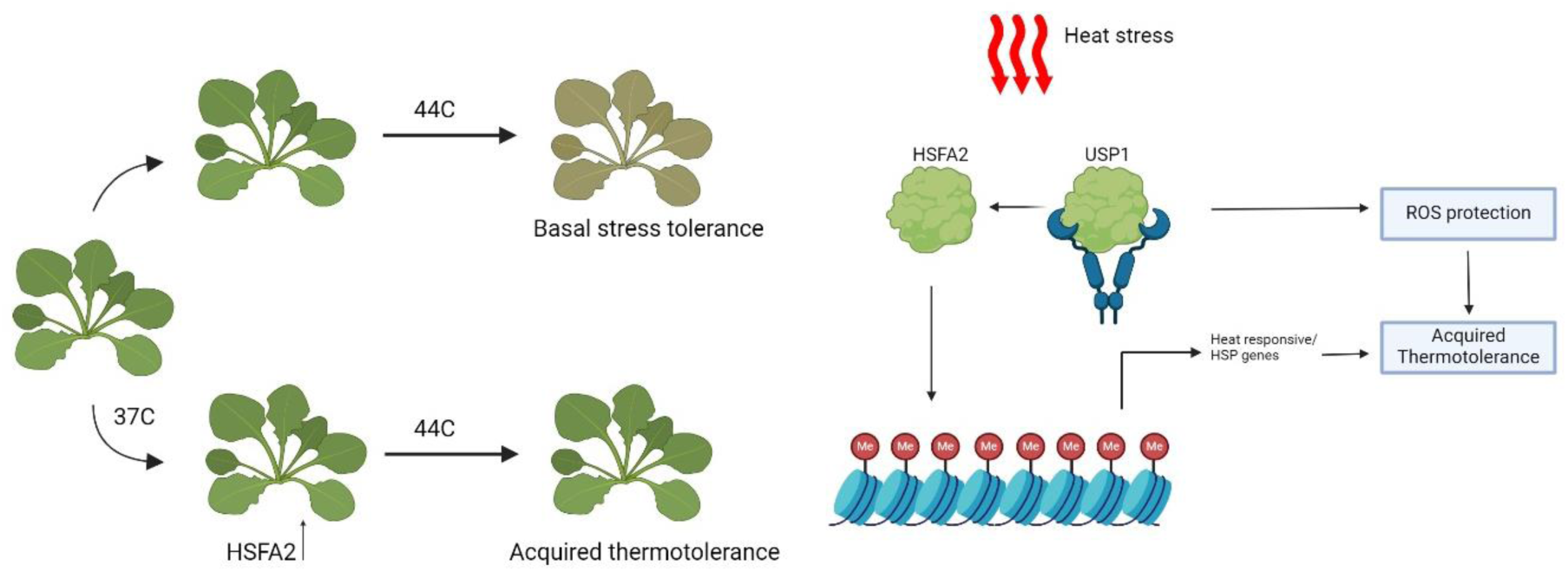
Proposed model of USP1, acting as a chaperone of HSFA2 during acquired thermotolerance in Arabidopsis. High temperature induces HSFA2 that regulates sustained histone H3K4 trimethylation at heat shock memory gene loci that activate expression *of HEAT SHOCK PROTEINs* (*HSPs*) leading to acquired thermotolerance. The images were generated using Biorender.com

## References

1. Byung-Hyun Lee and Sung-Hye Won and Hyo-Shin Lee and Mitsue Miyao and Won-Il Chung and In-Jung Kim and Jinki, J. (2000) ’Expression of the chloroplast-localized small heat shock protein by oxidative stress in rice’, Gene, 245(2), pp. 283–290.

2. Campbell, P. K. E., Huemmrich, K. F., Middleton, E. M., Ward, L. A., Julitta, T., Daughtry, C. S. T., Burkart, A., Russ, A. L. and Kustas, W. P. (2019) ’Diurnal and Seasonal Variations in Chlorophyll Fluorescence Associated with Photosynthesis at Leaf and Canopy Scales’, Remote Sensing, 11(5), pp. 488.

3. Cendrero-Mateo, M. P., Moran, M. S., Papuga, S. A., Thorp, K. R., Alonso, L., Moreno, J., Ponce-Campos, G., Rascher, U. and Wang, G. (2015) ’Plant chlorophyll fluorescence: active and passive measurements at canopy and leaf scales with different nitrogen treatments’, Journal of Experimental Botany, 67(1), pp. 275–286.

4. Charng, Y. Y., Liu, H. C., Liu, N. Y., Chi, W. T., Wang, C. N., Chang, S. H. and Wang, T. T. (2007) ’A heat-inducible transcription factor, HsfA2, is required for extension of acquired thermotolerance in Arabidopsis’, Plant Physiol, 143(1), pp. 251–62.

5. Chi, Y. H., Paeng, S. K., Kim, M. J., Hwang, G. Y., Melencion, S. M., Oh, H. T. and Lee, S. Y. (2013) ’Redox-dependent functional switching of plant proteins accompanying with their structural changes’, Front Plant Sci, 4, pp. 277.

6. Choudhury, F. K., Rivero, R. M., Blumwald, E. and Mittler, R. (2017) ’Reactive oxygen species, abiotic stress and stress combination’, The Plant Journal, 90(5), pp. 856–867.

7. Chuang, M. H., Wu, M. S., Lo, W. L., Lin, J. T., Wong, C. H. and Chiou, S. H. (2006) ’The antioxidant protein alkylhydroperoxide reductase of Helicobacter pylori switches from a peroxide reductase to a molecular chaperone function’, Proc Natl Acad Sci U S A, 103(8), pp. 2552–7.

8. 8. Daudi, A. and O’Brien, J. A. (2012) ’Detection of Hydrogen Peroxide by DAB Staining in Arabidopsis Leaves’, Bio Protoc, 2(18).

9. Dennis, G., Jr., Sherman, B. T., Hosack, D. A., Yang, J., Gao, W., Lane, H. C. and Lempicki, R. A. (2003) ’DAVID: Database for Annotation, Visualization, and Integrated Discovery’, Genome Biol, 4(5), pp. P3.

10. Du, Z., Zhou, X., Ling, Y., Zhang, Z. and Su, Z. (2010) ’agriGO: a GO analysis toolkit for the agricultural community’, Nucleic Acids Res, 38(Web Server issue), pp. W64–70.

11. El-Sappah, A. H., Rather, S. A., Wani, S. H., Elrys, A. S., Bilal, M., Huang, Q., Dar, Z. A., Elashtokhy, M. M. A., Soaud, N., Koul, M., Mir, R. R., Yan, K., Li, J., El-Tarabily, K. A. and Abbas, M. (2022) ’Heat Stress-Mediated Constraints in Maize (Zea mays) Production: Challenges and Solutions’, Frontiers in Plant Science, 13.

12. Fan, M., Gao, S., Yang, Y., Yang, S., Wang, H. and Shi, L. (2024) ’Genome-wide identification and expression analysis of the universal stress protein (USP) gene family in Arabidopsis thaliana, Zea mays, and Oryza sativa’, Genetica, 152(2), pp. 119–132.

13. Gill, S. S. and Tuteja, N. (2010) ’Reactive oxygen species and antioxidant machinery in abiotic stress tolerance in crop plants’, Plant Physiol Biochem, 48(12), pp. 909–30.

14. Gurley, W. B. (2000) ’HSP101: a key component for the acquisition of thermotolerance in plants’, Plant Cell, 12(4), pp. 457–60.

15. Hong, S. W. and Vierling, E. (2000) ’Mutants of Arabidopsis thaliana defective in the acquisition of tolerance to high temperature stress’, Proc Natl Acad Sci U S A, 97(8), pp. 4392–7.

16. Hong, S. W. and Vierling, E. (2001) ’Hsp101 is necessary for heat tolerance but dispensable for development and germination in the absence of stress’, Plant J, 27(1), pp. 25–35.

17. Hunseung Kang and Su Jung Park and Kyung Jin, K. (2013) ’Plant RNA chaperones in stress response’, Trends in Plant Science, 18(2), pp. 100–106.

18. Jagtap, A. B., Yadav, I. S., Vikal, Y., Praba, U. P., Kaur, N., Gill, A. S. and Johal, G. S. (2023) ’Transcriptional dynamics of maize leaves, pollens and ovules to gain insights into heat stress-related responses’, Frontiers in Plant Science, 14.

19. Jung, Y. J., Melencion, S. M. B., Lee, E. S., Park, J. H., Alinapon, C. V., Oh, H. T., Yun, D.-J., Chi, Y. H. and Lee, S. Y. (2015) ’Universal Stress Protein Exhibits a Redox-Dependent Chaperone Function in Arabidopsis and Enhances Plant Tolerance to Heat Shock and Oxidative Stress’, Frontiers in Plant Science, 6.

20. Kerk, D., Bulgrien, J., Smith, D. W. and Gribskov, M. (2003) ’Arabidopsis proteins containing similarity to the universal stress protein domain of bacteria’, Plant Physiol, 131(3), pp. 1209–19.

21. Kirti Shekhawat and Marilia Almeida-Trapp and Gabriel, X. G.-R. a. H. H. (2022) ’Beat the heat: plant- and microbe-mediated strategies for crop thermotolerance’, Trends in Plant Science, 27(8), pp. 802–813.

22. Larkindale, J., Hall, J. D., Knight, M. R. and Vierling, E. (2005) ’Heat stress phenotypes of Arabidopsis mutants implicate multiple signaling pathways in the acquisition of thermotolerance’, Plant Physiol, 138(2), pp. 882–97.

23. Larkindale, J. and Huang, B. (2004) ’Thermotolerance and antioxidant systems in Agrostis stolonifera: involvement of salicylic acid, abscisic acid, calcium, hydrogen peroxide, and ethylene’, J Plant Physiol, 161(4), pp. 405–13.

24. Lee, D. G., Ahsan, N., Lee, S. H., Kang, K. Y., Bahk, J. D., Lee, I. J. and Lee, B. H. (2007) ’A proteomic approach in analyzing heat-responsive proteins in rice leaves’, Proteomics, 7(18), pp. 3369–83.

25. Li, H., Li, H., Lv, Y., Wang, Y., Wang, Z., Xin, C., Liu, S., Zhu, X., Song, F. and Li, X. (2020) ’Salt Priming Protects Photosynthetic Electron Transport against Low-Temperature-Induced Damage in Wheat’, Sensors, 20(1), pp. 62.

26. Li, S., Fu, Q., Chen, L., Huang, W. and Yu, D. ‘Arabidopsis thaliana WRKY25, WRKY26, and WRKY33 coordinate induction of plant thermotolerance’.

27 Li, S., Fu, Q., Huang, W. and Yu, D. ‘Functional analysis of an Arabidopsis transcription factor WRKY25 in heat stress’.

28. Lin, M. Y., Chai, K. H., Ko, S. S., Kuang, L. Y., Lur, H. S. and Charng, Y. Y. (2014) ’A positive feedback loop between HEAT SHOCK PROTEIN101 and HEAT STRESS-ASSOCIATED 32-KD PROTEIN modulates long-term acquired thermotolerance illustrating diverse heat stress responses in rice varieties’, Plant Physiol, 164(4), pp. 2045–53.

29. Liu, H.-c., Lämke, J., Lin, S.-y., Hung, M.-J., Liu, K.-M., Charng, Y.-y. and Bäurle, I. (2018a) ’Distinct heat shock factors and chromatin modifications mediate the organ-autonomous transcriptional memory of heat stress’, The Plant Journal, 95(3), pp. 401–413.

30. Liu, H. C., Lämke, J., Lin, S. Y., Hung, M. J., Liu, K. M., Charng, Y. Y. and Bäurle, I. (2018b) ’Distinct heat shock factors and chromatin modifications mediate the organ-autonomous transcriptional memory of heat stress’, Plant J, 95(3), pp. 401–413.

31. Liu, Y. H., Tseng, T. S., Wu, C. R., Cho, S. T., Kuo, C. H., Huang, X. J., Cheng, J. Y., Hsu, K. H., Lin, K. F., Liu, C. C. and Yeh, C. H. (2023) ’Rice OsHsp16.9A interacts with OsHsp101 to confer thermotolerance’, Plant Sci, 330, pp. 111634.

32. Loukehaich, R., Wang, T., Ouyang, B., Ziaf, K., Li, H., Zhang, J., Lu, Y. and Ye, Z. (2012) ’SpUSP, an annexin-interacting universal stress protein, enhances drought tolerance in tomato’, Journal of Experimental Botany, 63(15), pp. 5593–5606.

33. Luo, D., Wu, Z., Bai, Q., Zhang, Y., Huang, M., Huang, Y. and Li, X. (2023) ’Universal Stress Proteins: From Gene to Function’, International Journal of Molecular Sciences, 24(5), pp. 4725.

34. Lämke, J., Brzezinka, K. and Bäurle, I. (2016) ’HSFA2 orchestrates transcriptional dynamics after heat stress in Arabidopsis thaliana’, Transcription, 7(4), pp. 111–4.

35. Malik, K., Matejtschuk, P., Thelwell, C. and Burns, C. J. (2013) ’Differential scanning fluorimetry: rapid screening of formulations that promote the stability of reference preparations’, J Pharm Biomed Anal, 77, pp. 163–6.

36. Marc, M., Cournol, M., Hanteville, S., Poisson, A.-S., Guillou, M.-C., Pelletier, S., Laurens, F., Tessier, C., Coureau, C., Renou, J.-P., Delaire, M. and Orsel, M. (2020) ’Pre-harvest climate and post-harvest acclimation to cold prevent from superficial scald development in Granny Smith apples’, Scientific Reports, 10(1), pp. 6180.

37. Melencion, S. M. B., Chi, Y. H., Pham, T. T., Paeng, S. K., Wi, S. D., Lee, C., Ryu, S. W., Koo, S.S. and Lee, S. Y. (2017) ’RNA Chaperone Function of a Universal Stress Protein in Arabidopsis Confers Enhanced Cold Stress Tolerance in Plants’, International Journal of Molecular Sciences, 18(12), pp. 2546.

38. Nishizawa, A., Yabuta, Y., Yoshida, E., Maruta, T., Yoshimura, K. and Shigeoka, S. (2006) ’Arabidopsis heat shock transcription factor A2 as a key regulator in response to several types of environmental stress’, Plant J, 48(4), pp. 535–47.

39. Nishizawa-Yokoi, A., Tainaka, H., Yoshida, E., Tamoi, M., Yabuta, Y. and Shigeoka, S. (2010) ’The 26S proteasome function and Hsp90 activity involved in the regulation of HsfA2 expression in response to oxidative stress’, Plant Cell Physiol, 51(3), pp. 486–96.

40. Park, J. W., Mieyal, J. J., Rhee, S. G. and Chock, P. B. (2009) ’Deglutathionylation of 2-Cys peroxiredoxin is specifically catalyzed by sulfiredoxin’, J Biol Chem, 284(35), pp. 23364–74.

41. 41. Schramm, F., Ganguli, A., Kiehlmann, E., Englich, G., Walch, D. and von Koskull-Döring, P. (2006) ’The heat stress transcription factor HsfA2 serves as a regulatory amplifier of a subset of genes in the heat stress response in Arabidopsis’, Plant Mol Biol, 60(5), pp. 759–72.

42. 42. Schramm, F., Larkindale, J., Kiehlmann, E., Ganguli, A., Englich, G., Vierling, E. and von Koskull-Döring, P. (2008) ’A cascade of transcription factor DREB2A and heat stress transcription factor HsfA3 regulates the heat stress response of Arabidopsis’, Plant J, 53(2), pp. 264–74.

43. Shikanai, T. (2014) ’Central role of cyclic electron transport around photosystem I in the regulation of photosynthesis’, Curr Opin Biotechnol, 26, pp. 25–30.

44. Sinha, P., Pazhamala, L. T., Singh, V. K., Saxena, R. K., Krishnamurthy, L., Azam, S., Khan, A. W. and Varshney, R. K. (2015) ’Identification and Validation of Selected Universal Stress Protein Domain Containing Drought-Responsive Genes in Pigeonpea (Cajanus cajan L.)’, Front Plant Sci, 6, pp. 1065.

45. Sousa, M. C. and McKay, D. B. (2001) ’Structure of the universal stress protein of Haemophilus influenzae’, Structure, 9(12), pp. 1135–41.

46. Tikkanen, M. and Aro, E. M. (2014) ’Integrative regulatory network of plant thylakoid energy transduction’, Trends Plant Sci, 19(1), pp. 10–7.

47 Trapnell, C., Roberts, A., Goff, L., Pertea, G., Kim, D., Kelley, D. R., Pimentel, H., Salzberg, S. L., Rinn, J. L. and Pachter, L. ‘Differential gene and transcript expression analysis of RNA-seq experiments with TopHat and Cufflinks’.

48. Wang, Z.-P., Xing, H.-L., Dong, L., Zhang, H.-Y., Han, C.-Y., Wang, X.-C. and Chen, Q.-J. ‘Egg cell-specific promoter-controlled CRISPR/Cas9 efficiently generates homozygous mutants for multiple target genes in Arabidopsis in a single generation’.

49. Wu, H.-C., Vignols, F. and Jinn, T.-L. (2019) ’Temperature Stress and Redox Homeostasis: The Synergistic Network of Redox and Chaperone System in Response to Stress in Plants’, in Asea, A.A.A. and Kaur, P. (eds.) Heat Shock Proteins in Signaling Pathways. Cham: Springer International Publishing, pp. 53–90.

50. Yamori, W. and Shikanai, T. (2016) ’Physiological Functions of Cyclic Electron Transport Around Photosystem I in Sustaining Photosynthesis and Plant Growth’, Annu Rev Plant Biol, 67, pp. 81–106.

51. Zbacnik, T. J., Holcomb, R. E., Katayama, D. S., Murphy, B. M., Payne, R. W., Coccaro, R. C., Evans, G. J., Matsuura, J. E., Henry, C. S. and Manning, M. C. (2017) ’Role of Buffers in Protein Formulations’, J Pharm Sci, 106(3), pp. 713–733.

52. Zhao, Y., Du, H., Wang, Y., Wang, H., Yang, S., Li, C., Chen, N., Yang, H., Zhang, Y., Zhu, Y., Yang, L. and Hu, X. (2021) ’The calcium-dependent protein kinase ZmCDPK7 functions in heat-stress tolerance in maize’, Journal of Integrative Plant Biology, 63(3), pp. 510–527.

